# Single-cell lineage trajectories and chromatin regulators that initialize antiviral CD8 T cell ontogeny

**DOI:** 10.1101/2021.08.11.456014

**Authors:** Huitian Diao, Runqiang Chen, Shanel M. Tsuda, Dapeng Wang, Megan A. Frederick, Jihye Kim, Pabalu P. Karunadharma, Gustavo Martinez, Adam J. Getzler, Clara Toma, Justin J. Milner, Thomas C. Venables, Donna M. Martin, Ananda W. Goldrath, Shane Crotty, Matthew E. Pipkin

## Abstract

Individual naive CD8 T cells activated in lymphoid organs differentiate into functionally diverse and anatomically distributed T cell phylogenies in response to intracellular microbes. During infections that resolve rapidly, including live viral vaccines^1^, distinct effector (T_EFF_) and memory (T_MEM_) cell populations develop that ensure long term immunity^2^. During chronic infections, responding cells progressively become dysfunctional and “exhaust”^3^. A diverse taxonomy of T_EFF_, T_MEM_ and exhausted (T_EX_) CD8 T cell populations is known, but the initial developmental basis of this phenotypic variation remains unclear^4–10^. Here, we defined single-cell trajectories and identified chromatin regulators that establish antiviral CD8 T cell heterogeneity using unsupervised analyses of single-cell RNA dynamics^11–13^ and an *in vivo* RNAi screen^14^. Activated naive cells differentiate linearly into uncommitted effector-memory progenitor (EMP) cells, which initially branch into an analogous manifold during either acute or chronic infection. Disparate RNA velocities in single EMP cells initiate divergence into stem, circulating, and tissue-resident memory lineages that generate diverse T_MEM_ and T_EX_ precursor states in specific developmental orders. Interleukin-2 receptor (IL-2R) signals are essential for formation and transcriptional heterogeneity of EMP cells, and promote trajectories toward T_EFF_ rather than T_EX_ states. Nucleosome remodelers *Smarca4* and *Chd7* differentially promote transcription that delineates divergent T_MEM_ lineages before cooperatively driving terminal T_EFF_ cell differentiation. Thus, the lineage architecture is established by specific chromatin regulators that stabilize diverging transcription in uncommitted progenitors.

## Main Text

To clarify the initial origins of T cell memory we generated longitudinal single-cell RNA-sequencing (scRNA-seq) datasets and used unsupervised methods to map single-cell trajectories that developed from naive CD8 T cells specific for *Lymphocytic choriomeningitis virus* (LCMV) early after infection of wildtype mice with strains that cause either an acute (Armstrong, LCMV_Arm_), or chronic (Clone 13, LCMV_Cl13_) infection (fig S1A–B) ^15^. On days and 8 post infection (pi), clonal TCR transgenic P14 cells that had been adoptively transferred and endogenous polyclonal MHC-I tetramter-reactive CD8 T cells (GP33^+^), which both recognize the LCMV epitope GP_33-41_ in MHC H-2D^b^, were isolated from the spleens of the same host mice, and libraries were generated in parallel with fresh naive CD8 T cells purified from separate P14 mice. Individual naive cells are recruited into the response over the first ~ 3 days following primary infection ^16^. Due to this asynchrony, we anticipated the time-series sampling would encompass multiple developmental states that compose initial antiviral CD8 T cell ontogeny.

Dimensionality reduction and Louvain cluster extraction of cells was performed on all samples simultaneously using similar numbers of randomly sampled cells from each experimental group to limit potential biological biases arising from changes in subset compositions at different time points, and the data are represented in the two-dimensional PAGA initialized force-directed (FA) embedding (**Fig 1A** and fig S1B) ^17,18^. Partition-based graph abstraction (PAGA) inferred single-cell paths based on correlations between clusters ^13^, which were numbered according to pseudotime (P0-P10) to define a potential developmental order (**Fig 1A–B,** Table S1). As expected, naive cells (P0) clustered apart from all activated cells, which separated into multiple clusters (P1-P10) (**Fig 1A** and fig S1C–D) and the pseudotime arrangement correctly predicted the actual time dependent emergence of cells in specific clusters (naive, vs days 5 and 8 pi) (**Fig 1C,** S1C-D and Table S1). The distribution of P14 and GP33^+^ CD8 T cells among the clusters was similar in LCMV_Arm_-infected mice (fig S1D and E). P14 cells isolated from LCMV_Arm_ and LCMV_Cl13_ infected hosts were distributed in similar clusters on day 5 pi, but contributed differentially to clusters on day 8 pi (**Fig 1C** and fig S2D and E). Cellular identities of the clusters were imputed using gene set enrichment analysis (GSEA) and “subtractive” gene expression signatures extracted from published bulk-RNA-seq data derived from phenotypically defined CD8 T cell subsets ^19,20^ (Table S2, https://github.com/TCellResearchTeam/T_cell_signature_Reference). All major T_EFF_, T_MEM_, T_EX_ and naive cell signatures were strongly enriched (p-val <0.05, NES) in at least one Louvain cluster (**Fig 1D** and fig S1 J-M), and demonstrated that cells corresponding to all mature T_EFF_, T_MEM_ and T_EX_ cell gene expression states arise within 8 days following acute or chronic LCMV infection. The PAGA-inferred paths between these states facilitated precisely defining developmentally regulated gene expression at the single-cell level, which extends previous longitudinal studies of bulk populations during acute infection ^21^.

**Figure 1.**
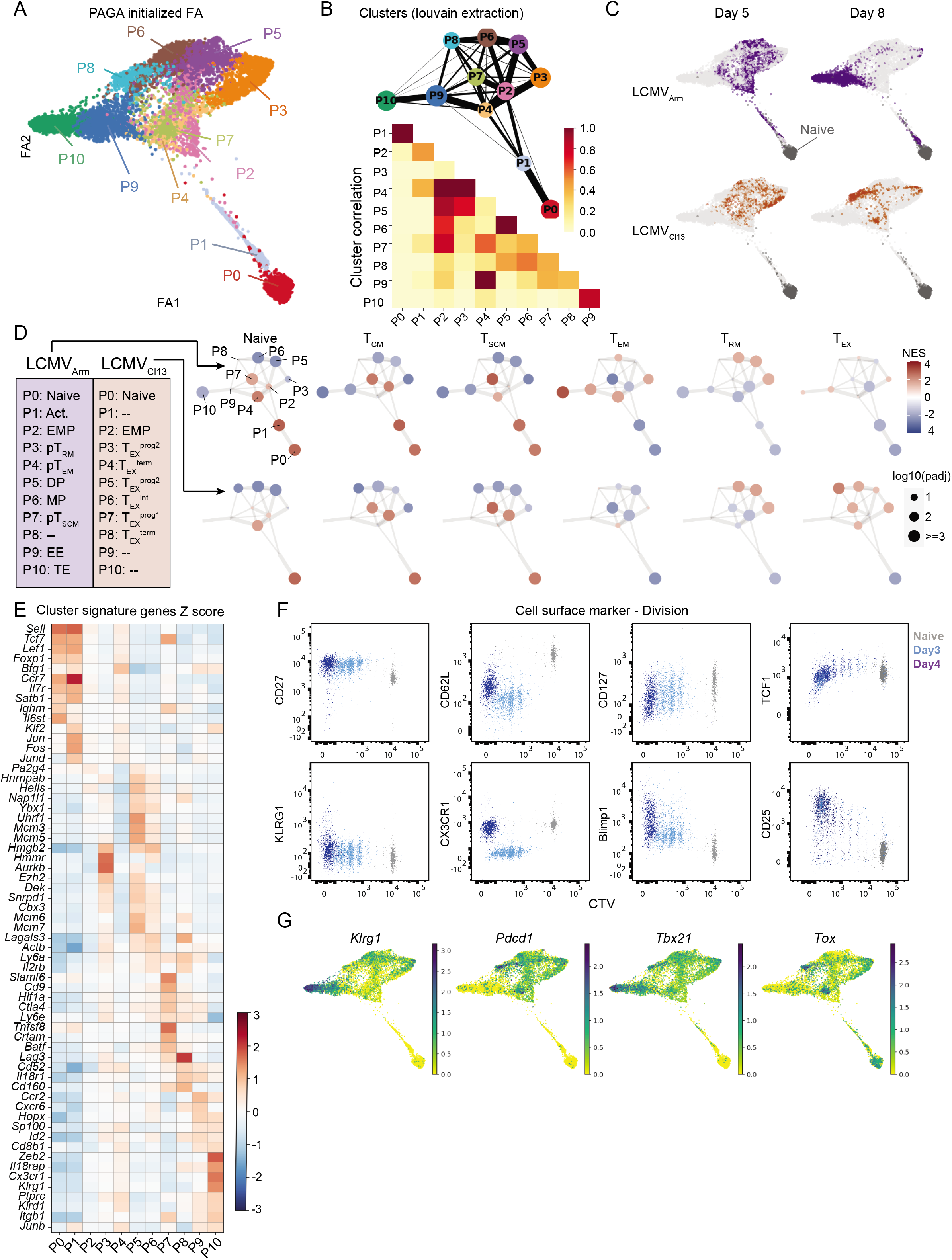
Naive CD8 T cells differentiate along a linear path into common effector and memory progenitor (EMP) cells. (A) PAGA initialized force-directed (FA) embedding based on individual cell gene expression profile, each dot represent one cell. PAGA connectivity analysis was performed on indicated clusters P0-P10. Clusters were extracted by Louvain method based on neighborhood graph. Neighborhood graph was calculated with UMAP algorithm. (B) Top: PAGA connectivity graph, each node represent one cluster, node sizes represent relative cell number of cluster, edge widths represent relative PAGA connectivity score. Bottom: heatmap of PAGA connectivity score between clusters. (C) PAGA initialized FA embedding, coloring cells based on origin: day5 /day8, naive (grey) / LCMV_Arm_ (purple) / LCMV_Cl13_ (orange). (D) GSEA of selected signatures for Louvain clusters, separating LCMV_Arm_ / LCMV_Cl13_. GSEA analysis performed on mean of normalized gene expression per cluster. NES represented by color, −log10(padj) represented by dot size. Labels of clusters inferred by GSEA result. (See also fig S1 J-M) (E) Mean of scaled expression, showing top signature genes of Louvain clusters. Signature genes were selected based on multiple differential analysis (Wilcoxon rank sum test, t-test, t-test over estimated variances) of cells within each cluster v.s. all others, with adjusted p-adj cutoff 0.05 (intersection of all tests) and absolute log2 fold change cutoff 1. Chromatin remodeling factors, transcription factors and surface proteins were selected from the genes that meet the statistical cutoffs for creating the signature gene lists. (See also Table S1). Top 10 signature genes ranked by t-test scores are represented in heatmap. (F) Expression of cell surface markers by CTV determined by flow cytometry. 50,000 P14 CD8 T cells were transferred per recipient mice which were given LCMV_Arm_. Naive cells (grey) and cells from day3 (blue) / day4 (purple) post infection are represented in plots. (G) Raw expression (logrithmized and normalized) of selected genes.

### Naive CD8 T cells differentiate along a linear path into common effector and memory progenitor (EMP) cells

The unsupervised approach clarified the initial developmental relationship of T_MEM_, T_EFF_, and T_EX_ cells in an ubiased fashion (**Fig 1B**). Naive cells (P0) were connected to cells in cluster P2, via activated intermediates (P1 cells, **Fig 1E** and Table S1). P2 cells were positively enriched with gene expression of recent TCR stimulation (48h Act up, p = 0.004, NES = 2.1). P4 cells were negatively enriched with this signature (fig S1K), and both GSEA and pseudotime indicated P4 cells were more developmentally advanced than those in P2, and were therefore downstream (fig S1J, Best clusters 2, 8 and 10). Thus, activated naive cells appeared to initially develop along a linear pathway into P2 cells.

Transcriptionally heterogeneous cell clusters on day 5 emerged from P2 cells, which strongly expressed *Il2ra* (encodes CD25/IL-2Rα, a subunit of the trimeric interleukin-2 receptor that initiates high-affinity IL-2 binding^22,23^) and were positively enriched with signatures of both central memory (T_CM_) and naive cells, but not those of mature effector memory (T_EM_), memory stem (T_SCM_), resident memory (T_RM_) or terminal effector (TE) cells (**Fig 1D** and fig S1L). P2 cells highly expressed a mixture of genes encoding TFs whose cognate motifs are enriched within *cis*-acting regions that gain *de novo* chromatin accessibility during primary TCR stimulation (*Runx3*, *Batf*, *Irf4, Prdm1*, *Klf2*), and that are essential for both T_EFF_ and T_MEM_ cell development^24,25^(fig S1G and H). In addition, genes encoding multiple regulatory factors whose expression is highly differential in established mature TE/T_EM_ (*Tbx21*, *Zeb2*, *Id2, Prdm1*), T_CM_/T_SCM_/T_EX_^prog1^ (*Tcf7*, *Zeb1*, *Bach2*, *Id3*), T_RM_ (*Hmmr, Aurkb, Prdm1*) and T_EX_^term^ (*Tox*, *Lag3, Cd160*) populations^26,27^ were coordinately expressed at intermediate levels in P2 cells (fig S1G and H). These “lineage-specific” genes were significantly upregulated or downregulated in cells from clusters at the distal tips of the paths (P10, P7, P3 and P8) compared to P2 cells (**Fig 1E**, fig S1G and H and Table S1), implying P2 cells promiscuously express regulatory factors that become progressively lineage-restricted. Flow cytometry confirmed that activated naive cells exhibited uniform behavior while undergoing extensive cell division, upregulating CD25 and maintaining expression of T_CM_ cell attributes (CD27 and Tcf1 expression), before developing phenotypic features of more mature T_EFF_ cells (e.g., high Blimp1-YFP and KLRG1 expression) (**Fig 1F**). These divergent subsets emerged near the final detectable cell division from cells highly expressing both CD25 and the naive and T_CM_-associated TF Tcf1 (fig S1I). Thus, naive cells initially differentiate in a linear path into cells that manifest multilineage gene expression, a hallmark of multipotency and cells undergoing lineage-choice ^28,29^. On this basis, we classified cells in cluster P2 as common effector/memory progenitor (EMP^P2^) cells.

### Disparate RNA velocities develop in individual EMP cells and initiate a branched manifold that establishes T_MEM_ and T_EX_ cell diversity

Strong connections of EMP^P2^ cells with clusters arranged immediately downstream implied the initial branchpoints of four developmental paths (**Fig 1B**). To define the trajectories of cells from each cluster in the PAGA-inferred architecture, their future states were modeled using RNA velocity^11,12^ (**Fig 2A**). Nascent RNA expression precedes accumulation of their mature mRNAs by several hours, and RNA velocity describes the rates at which cells are transitioning into new states based on the gene-wise ratios in expression of nascent (i.e. unspliced) to mature (i.e., spliced) mRNAs genome-wide. Streamline plots after UMAP embedding of all samples from each infection depicted transition probability data derived from the grid average RNA velocities between single-cell clusters (**Fig 2A** and fig S2A), and defined future cell states in the lineage architecture (**Fig 2B** and **2C** fig S2A). Strongly divergent RNA velocities in P2 and P5 cells confirmed they were developmental roots in each infectious context (fig S2B). The RNA velocities of signature genes associated with multiple distinct CD8 T cell states (e.g., naive and T_SCM_ cells: *Id3, Tcf7* and *Sell*; and TE cells: *Prdm1*, *Id2*, *Tbx21* and *Zeb2*) were all positive in EMP^P2^ cells during LCMV_Arm_ infection (**Fig 2E**, Table S5). This indicates that multilineage gene transcription in rapidly dividing EMP^P2^ cells establishes the transition potential into diverse future cell states, prior to developmental branching and mature lineage-specific mRNA expression. Conversely, cells in cluster P10 during LCMV_Arm_ infection, and clusters P7 and P8 during LCMV_Cl13_ infection, lacked RNA velocities into other clusters indicating they were terminal states in the analysis (fig S2C).

**Figure 2.**
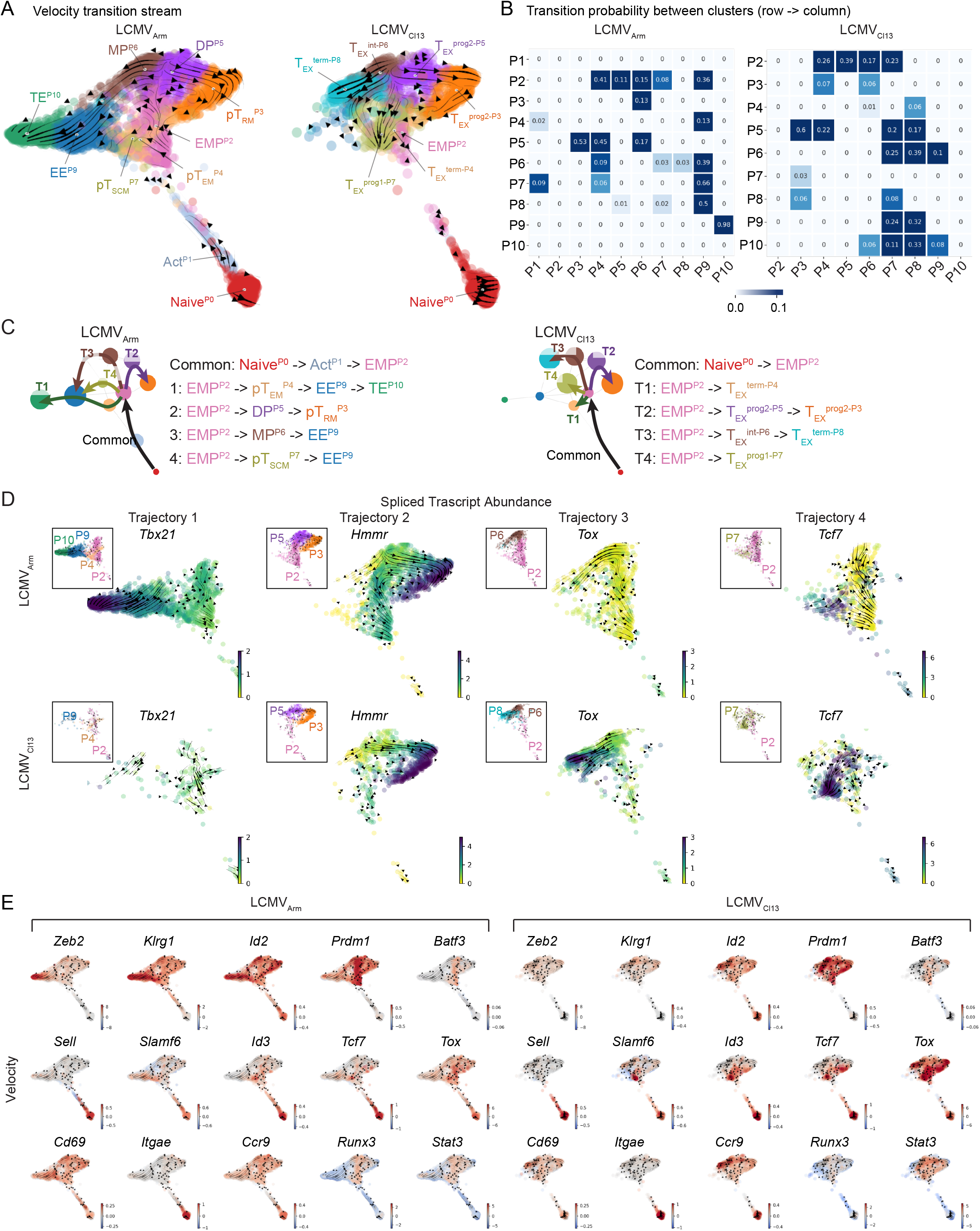
Disparate RNA velocities develop in individual EMP cells and initiate a branched manifold that establishes T_MEM_ and T_EX_ cell diversity. (A) Stream plot of velocity embedded on PAGA initialized single cell projection, separating LCMV_Arm_ / LCMV_Cl13_ (including naive cells in both conditions). (B) Transition probability heatmap between clusters estimated by scVelo, separating LCMV_Arm_ / LCMV_Cl13_. Color represent transition probability from row cluster to column cluster. (C) Inferred LCMV_Arm_ / LCMV_Cl13_ developmental rajectory by PAGA connectivity analysis, scVelo transition probability, pseudo-time (see Table S1) and real time. (D) Spliced transcript abundance (Ms) of representative genes for each of the four inferred trajectory in LCMV_Arm_ / LCMV_Cl13_. (E) Single cell velocity of selected driver genes, separating LCMV_Arm_ / LCMV_Cl13_. Potential driver genes were identified by combining top likelihood genes from analysis of all cells or multiple pairs of clusters with transitioning potential. Transcription factors, chromatin remodelers and surface receptors were selected within the identified likelihood genes (See Table S3).

Trajectory 1 (T1: P2->P4->P9->P10) only formed in LCMV_Arm_-infected mice and defined formation of cells enriched with signatures of bulk T_EM_ cells (effector memory precursors, pT_EM_^P4^), classical KLRG1^lo^ CD127^lo^ early effector cells (EE ^P9^) and KLRG1^hi^ CD127^lo^ terminal effector (TE) cells (TE^P10^) (**Fig 2C–D**, fig S1K and M). RNA velocity-derived transition probabilities indicated most EE^P9^ cells proceed toward TE^P10^ cells (**Fig 2B**). Positive *Tbx21* RNA velocity was sustained and *Zeb2* velocity accelerated in EE^P9^ cells, followed by increased expression of mature *Tbx21* and *Zeb2* mRNAs in TE^P10^ cells, whose velocities continued to increase in TE^P10^ cells^30–32^(**Figs 2D-E** and fig S2D and Table S5). Thus, the initial developmental order of T_CIRC_ cells during acute LCMV infection is T_CM_ to T_EM_ to TE cells.

Trajectory 2 (T2: P2>P5>P3) is likely a main source of T_RM_ precursor cells (pT_RM_^P3^) during LCMV_Arm_ infection, and exhausted progenitor 2 (T_EX_^prog2-P3^) cells^33^ during LCMV_Cl13_ infection. In LCMV_Arm_-infected hosts, P5 cells were positively enriched with signatures of bulk KLRG1^hi^CD127^hi^ “double positive” (DP) effector and T_RM_ cells, and were classified as DP^P5^ cells (**Fig 1D** and fig S1K). DP^P5^ cells diverged into pT_RM_^P3^ cells, pT_EM_^P4^ cells, and cells enriched with the bulk signature of classical KLRG1^lo^CD127^hi^ memory precursor (MP) cells (MP^P6^) (**Figs 1B and 2A-C** and fig S1K–L). Runx3-dependent gene expression was positively enriched in DP^P5^ and MP^P6^ (fig S1K and S2F), consistent with the requirement of Runx3 for development of these cell states^25,34^, and divergence into pT_RM_^P3^ cells correlated with increased expression of the reprsentative T_RM_ signature gene *Hmmr* (**Fig 2B–D**, and fig S1H). In LCMV_Cl13_-infected hosts P5 cells were also enriched with signatures of DP and T_RM_ cells, however, they were classified as T_EX_^prog2-P5^ cells because they were positively enriched with the signatures of both T_EX_^prog2^ and T_EX_ cells (**Fig 1D** and fig S1L–M). In addition, cluster P3 cells were enriched with both T_RM_ and T_EX_^prog2^ signatures, and were classified as T_EX_^prog2-P3^ cells, but were distinct from T_EX_^prog2-P5^ cells because they lacked TCR-stimulated gene expression (fig S1K–L). The T_EX_ signature was not strongly enriched in either DP^P5^ or pT_RM_^P3^ cells from LCMV_Arm_-infected mice, which confirmed their distinction from homologous cells during chronic infection which were and that strongly expressed *Tox*^35–37^ (**Figs 1D and 2E**, and fig S1M).

Trajectory 3 (T3: P2>P6>P8) explained initial development of intermediately exhausted (T_EX_^int^) and terminally exhausted cells (T_EX_^term^) during LCMV_Cl13_ infection, and formation of classical MP cells during LCMV_Arm_ infection (**Fig 2C**). In LCMV_Arm_-infected hosts, MP^P6^ cells were enriched with MP and EE cell signatures and had transition potential into EE^P9^ cells, providing an alternative conduit into the T1 trajectory (**Fig 2A–C,** fig S1K–M), and demonstrated classical MP cells are most likely distinct from precursors of T_SCM_ (see below). P6 cells from LCMV_Cl13_-infected were classified as T_EX_^int-P6^ cells because they manifested signatures of both intermediately exhausted (T_EX_^int^) and T_EX_ cells, and flowed directly into cluster P8 cells (**Fig 2A–D**), which were designated T_EX_^term-P8^ cells because they were strongly enriched with the T_EX_^term^ cell signature (**Fig 1D,** fig S1M) and highly expressed *Tox, Pdcd1* and *Lag3* (**Fig 1E and G**). Thus, T_EX_^int^ cells are related to classical MP cells, but exhibit altered transition probabilities into terminal states and fail to establish the T_CIRC_ lineage during LCMV_Cl13_ infection (**Fig 2B–D** and fig S2C).

Trajectory 4 (T4: P2> P7) defined formation of precursors of T_SCM_-like cells during acute (T_SCM_^P7^) or chronic (T_EX_^prog1-P7^) LCMV infection (**Fig 2C**). T_SCM_^P7^ cells from LCMV_Arm_- infected hosts were positively enriched with T_SCM_ and T_CM_ cell signatures, and were strongly connected to pT_EM_^P4^ and EE ^P9^ cells (**Fig 1D** and fig S1J–K). Both T_SCM_^P7^ and pT_EM_^P4^ cells exhibited strong *Tbx21* RNA velocity, suggesting they both manifest transition potential into *Tbx21* expressing states. Consistent with this, cells in both clusters exhibited strong transition probabilities into EE^P9^ cells, indicating that T_SCM_ precursors during LCMV_Arm_ infection are poised with T_EFF_ cell potential. In contrast, although T_EX_^prog1-P7^ cells from LCMV_Cl13_-infected hosts were positively enriched with the T_SCM_ signature, they exhibited reduced *Tbx21* RNA velocity (Table S5), and lacked transition probability into EE^P9^ cells (**Fig 2B**). In addition, T_EX_^prog1-P7^ cells appeared to derive from EMP^P2-Cl13^ cells initially, but their strong accumulation by day 8 pi correlated with retrograde transition potentials from all downstream T_EX_ cell clusters except T_EX_^prog2-P3^ cells, whereas during LCMV_Arm_ infection, T_SCM_^P7^ cells derived predominantly from EMP^P2^ cells (**Fig 2B–D**). Moreover, T_EX_^prog1-P7^ cells more strongly induced *Tcf7* (**Fig 2D**), highly expressed *Lag3*, *Pdcd1*, *Tox*, *Tox2* and *Bcl6*, were positively enriched with the specific T_EX_ cell signature and were enriched with terminal states (**Fig 1D** and figs S1H, M and S2C). Thus, the precursors of T_SCM_ cell states during acute and chronic infections have different origins and distinct developmental potentials.

### IL-2R-dependent transcription establishes EMP cells and transcriptional heterogeneity

The most dynamic genes drive the RNA velocity vector field ^11,12^. Those encoding TFs, CRFs, and surface receptors (SRs) during transitions between clusters were identified as potential drivers that compose the antiviral CD8 T cell architecture (**Fig 2C** and fig S2D–E, Table S4). Strong *Il2ra* dynamics and its transiently high expression in EMP^P2^ cells indicated IL-2R-dependent signals might promote multilineage potential (**Fig 3A and B**). To define the role of *Il2ra* functionally, single cell trajectories were inferred after scRNA-seq analysis of wildtype and *Il2ra*-deficient P14 (P14 *Il2ra*^−/−^) CD8 T cells 6 days after LCMV_Arm_ infection (**Fig 3C and D,** as described fig S1B). This second analysis confirmed the original lineage architecture during LCMV_Arm_ infection (**Fig 3C** and refer to **Fig 1A**). Louvain clusters from the *Il2ra* analysis (designated “Exp*-Il2ra*”) closely matched most clusters identified on days 5 and 8 pi in the original analyses (**Fig 3D** and refer to **Fig 1**). PAGA-inferred connectivity and RNA velocities implied analogous intracluster transition probabilities (**Fig 3C and E**), and analogous single cell behavior of wildtype P14 cells in both the original and Exp*-Il2ra* analyses emphasized that the inferred lineage architecture is biologically robust.

**Figure 3:**
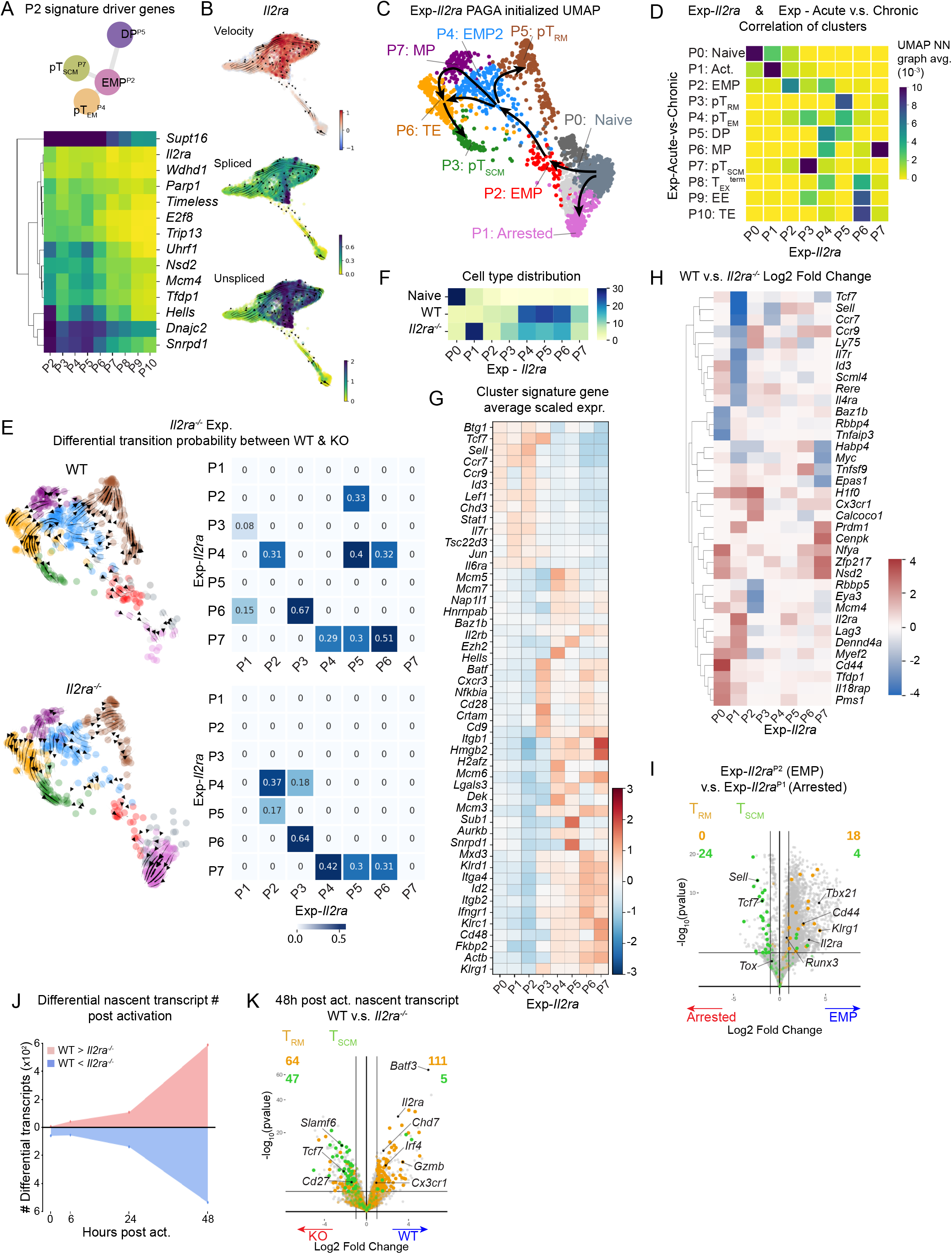
IL-2R-dependent transcription establishes EMP cells and transcriptional heterogeneity. (A) Heatmap of spliced transcript abundance of EMP^P2^ signature driver genes. EMP^P2^ common driver genes were defined as intersection between driver genes with likelihood > 0.25 in: all cells, P2-P4, P2-P5 and P2-P7 (Table S3) P2 signature driver genes were defined as intersection between P2 signature genes (described in **Fig 1E** and Table S1) and EMP^P2^ common driver genes. (B) Single cell normalized *Il2ra* transcript abundance, spliced (Ms) / unspliced (Mu). (C) PAGA initialized FA embedding based on single cell cell gene expression profile for Exp-*Il2ra*. PAGA connectivity analysis was performed on indicated clusters. Clusters were extracted from UMAP neighbor graph by Louvain method. (D) Correlation of clusters between acute v.s. chronic single cell experiment and Exp-*Il2ra*. Correlation represented by mean UMAP nearest neighbor graph scores between clusters from two experiments. UMAP projection of single cells from two experiments generated by Harmony integrated normalized count matrix from both experiment. (E) Transition probability between clusters in activated WT and *Il2ra*^−/−^ cells. Left: all single cells from WT / *Il2ra*^−/−^ samples in PAGA projection. Right: transition probability heatmap. (F) Percentage distribution in Louvain clusters for each cell type. Total percentage in all Louvain clusters add up to 100% for each cell type. (G) Signature gene heatmap for clusters (calculated for all cells, including naive, WT and *Il2ra*^−/−^). Method as described in **Fig 1E**. (H) Heatmap of log2 fold change of WT versus *Il2ra*^−/−^ for selected differential genes. Genes were selected from cluster signature gene list (described in **Fig 2G**). The genes with minimum expression of 0.0015 in at least one cluster and with absolute log2fc >= 2 in at least one cluster (WT versus *Il2ra*^−/−^) were used. (I) Differential analysis of transcript abundance comparing Exp-*Il2ra*^P1^ and Exp-*Il2ra*^P2^. X-axis represents log2 fold change, and the y-axis represents −log10(pvalue). The horizontal line indicates pval = 0.05. The vertical lines indicates absolute log2fc = 1. T_RM_ and T_SCM_ signature genes are highlighted in yellow and green respectively. (J) Number of differential nascent transcripts (between WT and *Il2ra*^−/−^) at different time points post activation. Differential nascent transcript is determined by DESeq2 (padj < 0.05). (K) Differential analysis volcano plot of WT versus *Il2ra*^−/−^ at 48 hours post activation. The x-axis represents log2 fold change, and y-axis represents −log10(pvalue). The horizontal line indicates pval = 0.05. The vertical lines indicates absolute log2fc = 1. T_RM_ and T_SCM_ signature genes are highlighted in yellow and green respectively.

P14 *Il2ra*^−/−^ CD8 T cells distributed within the trajectories aberrantly compared to WT P14 cells. P14 *Il2ra*^−/−^ cells almost entirely composed cluster Exp*-Il2ra*^P1^ compared to wildtype P14 cells indicating they arrested before transition into EMP^P2^ cells (Exp*-Il2ra*^P2^) (**Fig 3F** and fig S3E, p-value 1.35 × 10^−27^). Exp*-Il2ra*^P1^ cells were activated (data not shown), but lacked RNA velocity into future states indicating they were terminal (fig S3D). Strong differenital expression between Exp*-Il2ra*^P1^ and Exp*-Il2ra*^P2^ cells confirmed *Il2ra* was essential for transition into EMP^P2^ cells (**Fig 3I**). In addition, differential expression between wildtype and *Il2ra*^−/−^ P14 cells in cluster Exp*-Il2ra*^P2^ confirmed IL-2 signaling was required for EMP cell formation (fig S3G). This required IL-2R-dependent transcription, because genes whose nascent RNA expression required *Il2ra* for upregulation after TCR stimulation (WT^48h^ > WT^naive^, padj < 0.05) were positively enriched with those upregulated as Exp*-Il2ra*^P1^ transitioned into Exp*-Il2ra*^P2^ cells (fig S3K, NES = 1.26, pvalue = 0.04), whereas genes that required *Il2ra* for downregulation (WT^48h^ > *Il2ra*^−/−,48h^, padj < 0.05) were positively enriched with those downregulated in this transition (fig S3K, NES = − 1.16, pvalue = 0.05). Thus, IL-2R-dependent transcription *in vivo* is essential for gene expression that drives activated naive cells to become EMP^P2^ cells.

Development beyond the EMP^P2^ cell state also required IL-2R signals. The transition probabilities of P14 *Il2ra*^−/−^ cells into other clusters were substantially altered (**Fig 3E**). *Il2ra*^−/−^ cells in cluster Exp*-Il2ra*^P2^ lacked transition potential (**Fig 3E**). Those from cluster Exp*-Il2ra*^P4^ did not manifest velocity into Exp*-Il2ra*^P5^ (T_RM_) cells, and those in Exp*-Il2ra*^P5^ were vectored backward into cluster Exp-*Il2ra*^P2^ (**Fig 3E**). Consistent with this, P14 *Il2ra*^−/−^ cells were depleted from cluster Exp*-Il2ra*^P5^ cells (pT_RM_^P3^ analog) (**Fig 3F** and fig S3E, p-value 0.057), and those that did accumulate in that cluster were negatively enriched with the T_RM_-signature compared to wildtype P14 Exp*-Il2ra*^P5^ cells (fig S3F). Thus, *Il2ra*^−/−^ P14 cells that bypassed the initial developmental block inefficiently formed putative T_RM_ precursors. In addition, *Il2ra*^−/−^ cells in cluster Exp*-Il2ra*^P4^ manifested retrograde vectors into Exp*-Il2ra*^P3^ (T_SCM_) cells, unlike wildtype P14 cells (**Fig 3E**), which correlated with increased P14 *Il2ra*^−/−^ cell representation in cluster Exp*-Il2ra*^P3^ (T_SCM_^P7^ analogs) (**Fig 3F** and fig S3E, p-value 0.00042). Furthermore, *Il2ra*-deficient cells inefficiently repressed T_SCM_ signature genes after TCR stimulation (fig S3J). These results demonstrate that IL-2R signals promote divergent transcription in EMP cells that establishes trajectories into branching T_MEM_ cell lineages.

The bias of EMP-like *Il2ra*^−/−^ cells toward T_SCM_-like states and their reduced contribution to other T_MEM_ precursor states during acute infection prompted examining IL-2R-regulated genes in EMP^P2^ cells from LCMV_Cl13_-infected mice. EMP^P2^ cells during LCMV_Cl13_ infection less highly expressed IL-2R-Stat5 induced genes (*Batf3*, *Ccr5*, *Gzmb*, *Chd7* see below) that promote formation of protective T_EFF_ and T_MEM_ cells (fig. S3L). The IL-2R-repressed gene signature was enriched with genes whose RNA velocity was greater in EMP^P2^ cells from LCMV_Cl13_-infected mice (fig S3M). Moreover, in LCMV_Cl13_ infection, T_EX_^prog1-P7^ cells upregulated the IL-2R-repressed gene signature compared to EMP^P2^ cells; whereas this signature was not upregulated as EMP^P2^ transitioned to T_SCM_^P7^ cells during LCMV_Arm_ infection (fig S3M). Thus, regulation of IL-2R-dependent genes appear to bias the future states of EMP^P2^ cells during acute and chronic infection.

### Differential utilization of CRFs establishes antiviral CD8 T cell heterogeneity

Chromatin structure in naive cells is remodeled during initial TCR and IL-2 stimulation and becomes stably differential in distinct T_EFF_, T_MEM_ and T_EX_ cell subsets^38^. To identify chromatin regulatory factors (CRFs) that might control diverging transcriptional programs during T_EFF_ and T_MEM_ cell formation we screened a library of retrovirally delivered microRNA-embeded short hairpin RNAs (shRNAmirs) targeting nearly all murine CRF genes using a pooled approach in P14 CD8 T cells responding to LCMV_Arm_ infection (fig S4A and B, **Fig 5A** and Table S7)^14^. Candidate CRFs were identified by sequencing DNA libraries amplified from integrated shRNAmir proviral sequences in FACS-purified CD8 T cell subsets and quantifying differential shRNAmir representation (RNAi-induced effects) (**fig S4B**, https://github.com/Yolanda-HT/HSAP). Genes with top RNAi-effects (25^th^ percentiles) affecting *in vivo* P14 cell accumulation (input vs output), maturation of KLRG1^lo^ CD127^lo^ cells into all other phenotypes (EE vs other), and the balance between TE and MP cells (KLRG1^hi^CD127^lo^ vs KLRG1^lo^CD127^hi^) were selected as potential candidates (**Fig 4B–E** and Table S7). Individual follow-up experiments confirmed specific phenotypes of several top candidates that were previously unknown, including *Prmt5*, *Carm1*, *Taf1, Mll1* (manuscript in preparation) and multiple genes encoding factors in Brg1-associated factor (BAF, mammalian SWI/SNF)^39^ and chromodomian helicase and DNA-binding (Chd) nucleosome remodeling complexes (**Fig 4B–D**). Thus, concerted activity of many CRFs differentially control formation of classically defined populations defined by KLRG1 and CD127 expression during acute viral infection.

**Figure 4:**
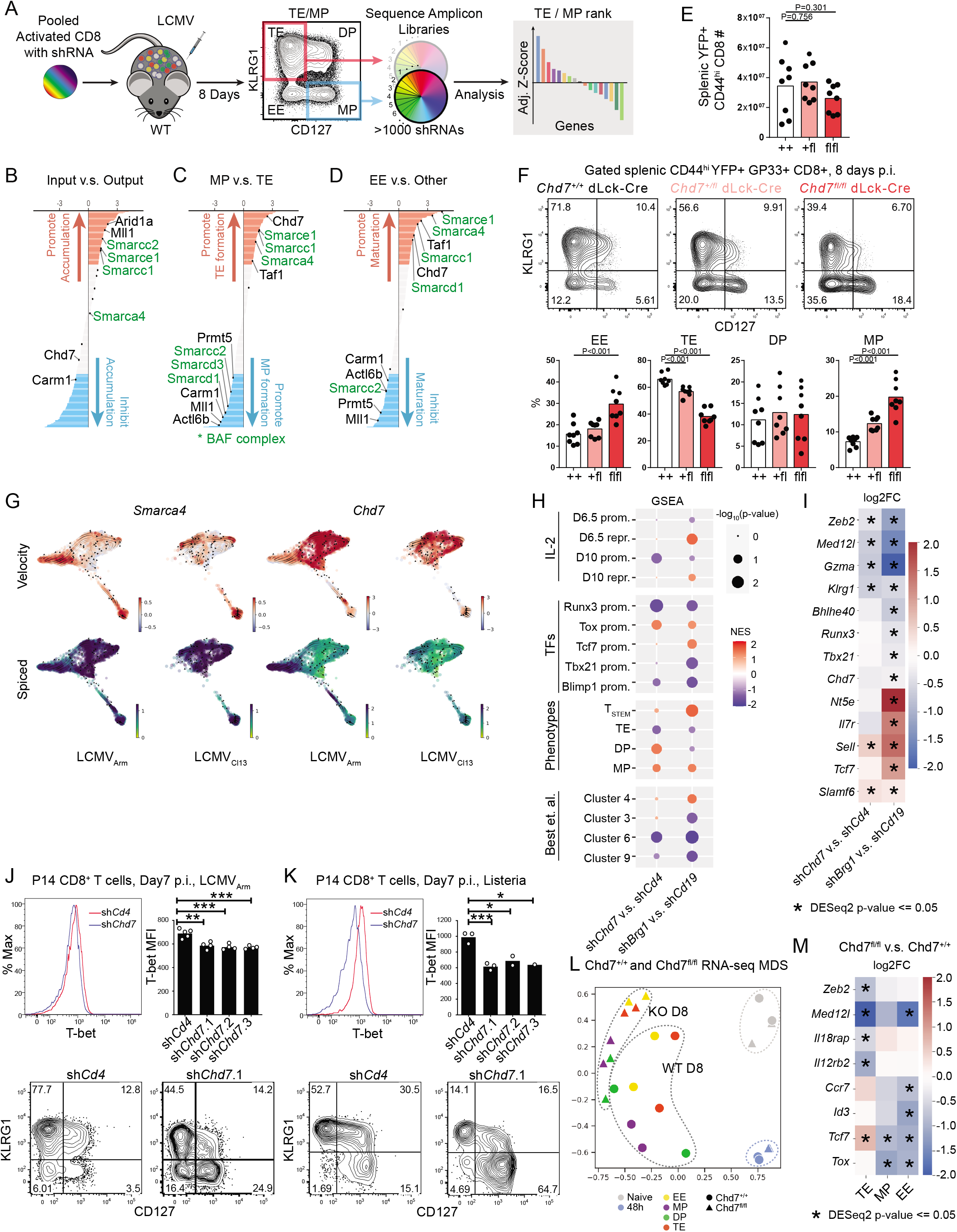
Differential utilization of CRFs establishes antiviral CD8 T cell heterogeneity. (A) Pooled RNAi screen and analysis simplified schematic (detailed schematic in fig S4A-B). (B-D) Ranked lists of adjusted RNA - effects for input versus output, MP versus TE, and EE versus others. Red / blue highlight genes that are top / bottom quarter in effect ranking. (E-F) Experiment characterizing endogeneous CD8^+^ T cells response to LCMV_Arm_ infection (day 8 pi), in mice of Chd7^fl/fl^, Chd7^+/fl^, Chd7^+/+^ genotypes. (E) Total activated CD8 T cell number in spleens. (F) Representative flow cytometry plots showing CD127 and KLRG1 staining, and summarized percentages of cells in each category. (G) Smarca4 and Chd7 velocity and spliced transcript abundance (Ms) in single cell projection, separating LCMV_Arm_ / LCMV_Cl13_. (H) GSEA of sh*Chd7* versus sh*Cd4* (control) and sh*Brg1* versus sh*Cd19* (control). Signatures: IL-2 regulated signatures / key CD8 transcription factor (TF) regulated signatures / CD8 phenotype signatures / Best et. al. longitudinal expression dynamic gene signatures. (I) Differential expression of selected genes. Heatmap showing log2 fold change of gene expression comparing sh*Chd7* versus sh*Cd4* (control) or sh*Brg1* versus sh*Cd19* (control). (J-K) Comparison of T-bet expression and phenotype between sh*Chd7* versus sh*Cd4* (control) during LCMV_Arm_ or Listeria infection. Top: representative and summarized T-bet MFI. Bottom: representative flow cytometry plot showing CD127 and KLRG1 staining. (L) Multidimensional scaling plot showing similarity / dis-similarity of Chd7^fl/fl^ versus Chd7^+/+^ cells in different stages: naive / 48h post activation / day8 sorted subsets. (M) Differential expression of selected genes. Heatmap showing log2 fold change of gene expression between Chd7^fl/fl^ versus Chd7^+/+^ in day8 sorted subsets (TE, MP, EE).

We pursued *Smarca4* (Brg1) and *Chd7* (Chd7), which encode ATPases of nucleosome remodeling complexes that have essential roles in multipotent neural crest cells^40^, human development and immune system function^41–43^, and thymic T cell development^44^. Their strong phenotype in the screen and their dynamics in the single-cell trajectory, both implied they could have essential functions in EMP cells. Depletion of either factor alone impaired formation of KLRG1^hi^ cells on day 5 pi, and increased the fractions of KLRG1^lo^ CD127^lo^ EE-like cells by day 8 pi, which suggested both factors might promote divergence into TE, DP or MP populations^4^ (**Fig 5B**). In addition, RNAi-effects in the primary screen indicated *Smarca4* and at least 4 additional BAF subunits (*Arid1a, Smarcc1*, *Smarce1*) were selectively required for TE cell formation (**Fig 4C**). Four other BAF-subunits (*Actl6b*, *Smarcc2*, *Smarcd1*, and *Smarcd3*) were preferentially required for MP formation (**Fig 4C**). Thus, distinct BAF-complex assemblies might differentially bias TE and MP cell development^39^. To further confirm the role of Chd7, engineered *Chd7* alleles^42^ were conditionally disrupted in mice using transgenic Cre expression in post-thymic T cells (referred to as *Chd7*^fl/fl^ dLck-Cre sfYFP). Similar to RNAi, *Chd7* gene-disruption impaired the frequency of KLRG1^hi^ cells 5 days after LCMV_Arm_ infection (fig S4C), and reduced formation of TE cells while increasing frequencies of EE- and MP-like cells 8 days after infection, without strongly affecting overall T_EFF_ cell numbers (**Fig 4E**). Thus, both BAF and Chd7 complexes are essential for the early phenotypic heterogeneity of T_EFF_ cells during acute infection.

### Smarca4 and Chd7 are required for transcriptional heterogeneity of EMP cells

Distinct RNA dynamics of *Smarca4* and *Chd7* in the trajectory implied sequential early functions during initial lineage divergence. Mature *Smarca4* mRNA expression was greatest in EMP^P2^ cells, before *Chd7* RNA velocity increased in EMP^P2^, DP^P5^ and MP^P6^ cells (**Fig 4G**). Mature *Chd7* transcripts were upregulated in EE^P9^ and TE^P10^ cells (**Fig 4G**) and in bulk KLRG1^hi^ cells on day 5 of LCMV_Arm_ infection^25^. *Chd7* transcription required *Il2ra* during TCR stimulation (**Fig 3K**). These data are consistent with Smarca4 and Chd7 functioning in an IL-2R-dependent transctiptional network. To examine this possibility, P14 cells depleted of either *Smarca4* or *Chd7* were analyzed by RNA-seq 5 days after LCMV_Arm_ infection. *Smarca4* was required to downregulate genes that are repressed by IL-2R (day 6), and to repress both the T_SCM_ and *Tcf7*-promoted gene expression signatures ^21,25,36,45–47^ (**Fig 4H**), whereas *Chd7* was required for activation of genes that require IL-2R for expression at later times (day 10)^46^ (**Fig 4H**). In addition, both factors were necessary for promoting gene expression activated by the TF Runx3 and repressing gene expression promoted by the TF Tox (**Fig 4H**). Thus, IL-2R-dependent transcriptional heterogeneity that develops in EMP cells early during acute infection required both Smarca4 and Chd7. In addition, the distinct requirements of each factor indicated they differentially promote transcription that drives divergence between T1 (pT_EM_), T2 (MP) and T4 (pT_SCM_) trajectories.

Smarca4 and Chd7 stabilized transcription that drives formation of the T1 trajectory. *Smarca4*-depleted P14 cells on day 5 pi expressed significantly less *Bhlhe40, Chd7, Gzma, Med12l, Runx3*, *Tbx21* and *Zeb2* (**Fig 4I**, *left*); *Chd7*-depleted cells expressed significantly less *Gzma, Il12rb2*, *Il18rap*, *Med12l* and *Zeb2*, while expression of *Bhlhe40, Batf3,* and *Tbx21* trended lower (**Fig 4I**, *right*). In P14 cells depleted of Chd7, T-bet protein expression and TE cell formation was more strongly impaired during LM_GP33_ infection than during LCMV_Arm_ infection, consistent with grossly impaired *Il12rb2* expression (**Fig 4I**, *right*), and the increased IL-12 concentrations during LM infection compared to LCMV infection^30,48^ (**Fig 6J-K** and data not shown). Complementation of Smarca4 or Chd7 depleted P14 cells with retrovirally expressed Tbx21 restored the normal pattern of TE and MP in each case (fig 4D and E), indicating they each promote T_EM_ and TE differentiation by ensuring *Tbx21* expression. However, enforced T-bet expression in the absence of *Chd7* did not resuce defective Gzmb expression, indicating that Chd7 is broadly required for cytolytic effector cell programming (data not shown). Because altered gene expression in the absence of Smarca4 and Chd7 is manifest on day 5 pi, prior to EE^P9^ cell formation, these results demonstrate that both CRFs are necessary to establish T_EFF_ gene expression prior to when cells with these phenotypes manifest.

### Chd7 is essential for T_MEM_ cell lineage branching

P14 cells depleted of Chd7 during LCMV_Arm_ infection were positively enriched with gene expression signatures of DP^P5^ and MP^P6^ cells on day 5 pi, which correlated with increased frequencies of EE and MP-phenotype cells in *Chd7*^fl/fl^ dLck-Cre sfYFP mice 8 days after LCMV_Arm_ infection (**Fig 4F**). Thus, cells lacking Chd7 appeared to arrest at the point where EMP^P2^ cells undergo branching into MP^P6^ and EE^P9^ cells, well before maturation of TE^P10^ cells, which brought into question whether they correctly stabilized the specific gene expression programs related to each of these flow cytometry phenotypes. To examine this directly, LCMV-specific CD8 T cells from *Chd7*^+/+^ and *Chd7^fl/fl^* dLck-Cre^+^ sfYFP^+^ mice that exhibited MP, EE, DP and TE cell phenotypes 8 days after LCMV_Arm_ infection were FACS-purified and analyzed using RNA-seq. Multidimensional scaling showed these populations from wildtype mice were separated from each other, whereas those from *Chd7*-deficient mice grouped (**Fig 4L**), indicating that gene expression states which diverged in wildtype T_EFF_ subsets did not strongly diverge in the *Chd7*-deficient populations. Consistent with this interpretation, pairwise analysis showed that compared to wildtype cells, *Chd7*-deficient TE-phenotypic cells less strongly expressed genes encoding factors characteristic of TE cells (*Zeb2*, *Med12l*, *Il18rap, Il12rb2*), and *Chd7*-deficient EE- or MP-phenotypic cells less highly expressed genes that promote MP cell development and formation of long-lived T_MEM_ cells (*Tcf7*, *Id3*, *Tox,* and *Ccr7*) (**Fig 4M**). These results demonstrate that Chd7 is necessary to stabilize divergent transcriptional programs that differentiates circulating T_MEM_ lineage branches and maturing cell states, and promotes terminal T_EFF_ maturation.

## Discussion

Our study resolves the initial stages of antiviral CD8 T cell ontogenesis. The architecture indicates naive cells differentiate into common EMP cells which diverge along distinct trajectories that develop gene expression states within the first week of acute or chronic viral infection that match all major T_EFF_, T_MEM_ and T_EX_ cell populations found at later times. Additional developmental paths to cells that were not sampled in this analysis could exist (e.g., cells from other tissues)^49,50^. Variable RNA velocities that develop in EMP cells indicates that diverging transcription initiates the branching trajectories before strong differential expression of lineage-specific regulators is established. IL-2R signals were required for EMP cell development and their transcriptional heterogeneity, and altered IL-2R-dependent transcription in EMP cells during chronic infection correlated with development of T_EX_ cell states. Because *Il2ra* is regulated by IL-2 stimulation and was dynamic during EMP cell formation and divergence, these results suggest variable IL-2 stimulation contributes to initial lineage bias of cells in the population.

We provide evidence that diverse T_MEM_ cell types most likely arise from divergent precursor cell states derived from distinct lineages early in the response. The nucleosome remodeler Smarca4 was necessary for gene expression that initially separates T_SCM_ and T_EM_ precursor states; Chd7 functioned later to mature classical MP cells; and, both factors cooperatively promoted TE cell differentiation. These results provide evidence that diverging transcriptional states in EMP cells are stabilized by specific CRFs, and implies that chromatin remodeling reinforces initial lineage biases and establishes the branching architecture. Thus, differential chromatin remodeling in divergent T_MEM_ cell precursor lineages might explain the preferential interconversion potentials of distinct mature T_MEM_ cell populations^1,51,52^, and resistance of T_EX_ cells to chromatin-level reprogramming^53,54^. The developmental architecture, stepwise transcriptional dynamics and CRF atlas described here suggests many factors with spatiotemporally resolved functions and might suggest strategies for engineering T_MEM_ CD8 T cell formation.

## Methods

### Mice

Wildtype 6-8 week old C57BL/6J mice were used as recipients for adoptive transfer experiments and were purchased from the Jackson Laboratory. P14 Thy1.1^+^ mice were a gift from Dr. Rafi Ahmed (Emory University). P14 Thy1.1^+^ Thy1.2^+^ mice were generated by crossing P14 Thy1.1^+^ mice with wildtype C57BL/6J mice. P14 Thy1.1^+^ *Il2ra*^−/−^ mice were generated by crossing *Il2ra*^−/−^ mice (purchased from Jackson Laboratory) with P14 Thy1.1^+^ mice. *Chd7*^fl/fl^ mice were a gift from Dr. Donna M. Martin (University of Michigan) ^42^, and were crossed with Rosa26-EYFP and dLck-Cre (maintained heterozygous) transgenic mice. All mice were maintained in specific-pathogen free facilities and used according to protocols approved by the Institutional Animal Care and Use Committee of TSRI-FL.

### T cell activation, adoptive transfer and infections

Naive P14 CD8^+^ T cells from wildtype mice were isolated by negative selection (EasySep™, Stemcell Technologies). Naive P14 Thy1.1^+^ *Il2ra*^−/−^ cells were isolated from 4-5 week old mice by depleting CD44^hi^ cells (Biolegend biotin anti-mouse/human CD44, clone IM7, 2μl per spleen). For *Il2ra*^−/−^ single cell experiment, cells were further sorted for CD44^lo^ with FACS. Purified naive P14 CD8^+^ T cells were resuspended in serum free DMEM and transferred by retro-orbital injection. For scRNA-seq experiments during acute and chronic LCMV infection, 2×10^4^ naive P14 CD8^+^ T cells were transferred per recipient mouse. For *Il2ra*^−/−^ single cell experiment, 2.4×10^5^ naive P14 CD8^+^ T cells were transferred per recipient mouse (Thy1.1^+^ *Il2ra*^−/−^ to Thy1.1^+^ Thy1.2^+^ ratio = 2:1). For retroviral transduction, purified naive CD8^+^ T cells were activated with anti-CD3 and anti-CD28, retrovirus was packaged and cells were transduced as described ^25^ with the following modifications. Naive CD8 T cells were activated for 16-18 hours, transduced for 4 hours with retroviral supernatants, and immediately transferred to naive 6-8 week old C57BL/6J hosts. 50,000 cells were transferred to each host and were infected with 2×10^5^ PFU of LCMV_Arm_, or 500,000 cells were transferred and hosts were administered IP injection of 1.5×10^5^ PFU of LCMV_Cl13_, or 1×10^4^ CFU of LM_GP33_. LCMV_Arm_, LCMV_Cl13_ and LM_GP33_ stocks were produced as described ^25^. Infections were administered ~1 hour after adoptive transfer of transduced T cells, or the following day(s) after naive cell transfer. Virus stocks were stored at −80°C and thawed immediately before dilution. IP injection of 2×10^5^ PFU of LCMV_Arm_ per mouse was used to initiate acute infection, and retroorbital injection of 2×10^6^ PFU LCMV_Cl13_ per mouse was used to initiate chronic infection.

### Flow cytometry analysis and Sorting for single cell sequencing

Single cell suspensions were prepared by disrupting spleen sections by pressing through 70μm cell strainer in DMEM with 10% FBS. The splenocyte suspensions or heparnized peripheral blood collections were pelleted and red blood cells were lysed using ACK buffer. Cells were resuspended in FACS buffer, stained for surface proteins and then fixed in 2-4% PFA and permeabilzed for intracellular staining. Anti-mouse CD4 (RM4-5), CD8 (53-6.7), CD44 (IM7), CD62L (MEL-14), CD25 (3C7), CD127 (A7R34), KLRG1 (2F1/KLRG1), CD27 (LG.3A10), TCF-1 (S33-966), CX3CR1 (SA011F11) and were purchased from Biolegend or BD Biosciences. Intracellular staining for TCF-1 was performed using the Foxp3 transcription factor staining kit (eBioscience). For analysis of cells on days 3 or 4 of infection, spleens were cut into 1-2mm pieces and digested with collagenase IV (100 U/mL, Worthington) and DNase I (10 μg/mL, Sigma) in complete T cell media for 10 min at 37°C on a nutator, then disrupted over a 70μm cell strainer. For FACS isolation, CD8^+^ T cells were initially enriched from total splenocyte preparations by negative selection with anti-CD4, anti-CD19, anti-B220 and anti-TER119 and magnetic streptavidin beads (Stemcell Technologies). Enriched cells were pre-stained with GP33-AF488 tetramer (NIH tetramer facility) to label endogenous LCMV-specific CD8^+^ T cells, followed by staining with CD8 (BV421), Thy1.1 (PE) and Thy1.2 (APC) to label donor P14 CD8^+^ T cells and the cells were sorted using a BD FACS Aria™ Fusion.

### *In vivo* Pooled RNAi Screen

The RNAi screen was performed and analyzed as shown in fig S4A–B and as previously described ^14^ with the following modifications and details. Naive P14 CD8 T cells were activated for 18 hours using plate bound anti-CD3 and soluble anti-CD28 and transduced for 4 hours in 96 well plates before cells from all wells were pooled. Immediately after pooling (~24hrs post activation) aliquots of 500,000 cells were transferred into multiple naive host mice that were rested for ~1 hour after receiving cells before inoculation with 1.5×10^5^ PFU LCMV_Cl13_ per mouse, which induces an acute infection in this setting^14^. Two entire biological replicates of the screen were performed and used for compuationatl analysis, and each replicate was screened in three pools. Each pool of shRNAmirs targeting CRFs was generated from ~500 shRNAmirs shRNAmirs which also included a common set of control shRNAmirs. Each pool was analyzed in 10 recipient mice to maintain 50-100-fold theoretical representation of each shRNAmir after engraftment. For input samples, ~8×10^5^ transduced cells were FACS-purified 48 hours after transduction. Eight days following infection, 3-8×10^5^ cells from each KLRG1/CD127 gate were recovered by FACS from the spleens of infected mice. Genomic DNA was purified from each sample and used as PCR template to generate libraries for high-throughput sequencing as described ^14^. Sequencing reads are mapped to library fasta file containing shRNA sequence information with custom blast pipeline. Raw read counts for each shRNA are normalized to total counts, and quantiles of each shRNA were calculated with negative binomial distribution. To calculate the effect size of shRNA in different cell population, the difference between quantiles in different cell population (quantile shift) was calculated for individual shNRAs. To sum up the effect of each gene, the quantile shift for all shRNAs towards each gene were converted to Z-scores, which were averaged and adjusted by p-value (to account for the consistency of effect) to generate the adjsusted Z-Scores for ranking (fig S4B).

### Nascent RNA-seq analysis

For nascent RNA-seq of *in vitro* activated CD8 T cells, chromatin-associated RNA was prepared and total RNA-seq libraries were prepared and sequenced as described ^25^. Pair end fastq reads were trimmed with “trim_galore --paired --length 24 --stringency 3”. Trimmed fastq reads were aligned to GRCm38 genome with bowtie2 with parameters “-p 16 -X 2000 --fr” ^57^. Forward and reverse strand reads were separated with samtools ^58^. Reads per transcript were counted with subread featureCounts ^59^. Counts from forward and reverse strands were merged with custom python script. Differential analysis was performed with DEseq2 ^60^.

### Single cell RNA-seq library generation, sequencing and analysis

To prepare barcoded scRNA-seq libriaries from multiple libraries, anti-mouse mouse MHC H-2 hashtag antibodies (Biolegend TotalSeq™) were used to label separate FACS-purified subsets which were washed, counted and pooled to final concentration of ~1,600 cells/ul. A total of ~50,000 cells were loaded into one lane of the single cell A chip kit (P/N 1000009). Single-cell gel beads in emulsion (GEM) were generated using the 10X Chromium single cell controller (10X Genomics, Pleasanton, CA). Single-cell GEM’s and sequencing libraries were generated using the Single-cell 3’ library and gel bead kit V2 (P/N 120267) according to manufacturer recommendations. The final library size distributions and amounts were assessed using bioanalyzer analysis and further quantified with the NEBNext library quantification kit (P/N E7630). The cDNA and HTO libraries were pooled 10:1 and sequenced to a depth of 50,000 reads per cell for the cDNA and 2,000 reads per cell for the HTO library. Libraries were sequenced on the Illumina NovaSeq with the following 10X read format; Read 1 25bp, index i7 8bp, and Read 2 98bp. Around 500 - 1500 million reads were generated per experiment, yielding 60-90% sequencing saturation and around 1500-3000 median genes per cell after alignment. For hashtag library sequences, 60-75% antibody reads were mapped to cells (usable), yielding around 3000-6000 usable antibody reads per cell.

Cellranger 3.0 was used for fastq generation (cellranger mkfastq) and counting (cellranger count). Standard quality filtering was performed with Scanpy to remove genes expressed in less than 3 cells, doublets or low read cells (with gene / count per cell cutoff), and cells with high mitochondria count percentage ^61^. Demultiplexing of Biolegend hashtags was performed with a custom python script to exclude doublets and dropouts. After quality filtering, reads were normalized to 10,000 per cell and logarithmized. Highly variable genes were identified with scanpy.pp.highly_variable_genes and selected. Count matrices were then regressed and scaled with scanpy.pp.regress_out and scanpy.pp.scale. Force-directed embedding was generated after PCA (scanpy.tl.pca) and UMAP based nearest neighbor analysis (scanpy.pp.neighbors). Louvain cluster extraction (scanpy.tl.louvain) was performed on force-directed embedding and the extracted clusters were analyzed with PAGA (scanpy.pl.paga) for cluster correlations ^18 13^. For velocity-based analysis, Velocyto was used to generate spliced / unspliced count matrix ^12^. The resulting matrix was processed in scVelo for velocity analysis ^11^. For correlation of independent scRNA-seq experiments, Harmonypy was used for batch effect removal on normalized and scaled count matrices of experiments ^62^. Dimensionality reduction was performed on batch effect removed matrices with PCA, followed by UMAP projection.

### ChIP-seq analysis

Raw fastq files of ChIP-seq experiments were downloaded from GEO with SRA-Tools fastq-dump. Trimming of fastq files were performed with trim_galore. Trimpped reads were aligned to GRCm38 genome with bowtie2 ^57^. Aligned sam file were sorted and filtered for PCR duplicates with samtools. Blacklisted regions were filtered with bedtools ^63,64^. Peaks were called with MACS2 and annotated with R package ChIPSeeker ^65,66^. All analytical codes for ChIP-seq are published on Github: https://github.com/TCellResearchTeam/T_Cell_ChIP. Visualization of ChIP data is accessible on <u>UCSC Track Hub</u>: T_cell_ATAC_ChIP_Pipkin.

### GSEA

GSEA was performed with R package clusterProfiler ^20^. GSEA signatures were downloaded or generated from published datasets available from GEO database. All signatures and description are available on https://github.com/TCellResearchTeam/T_cell_signature_Reference.

## Acknowledgments

This work was supported by Scripps Florida via the State of Florida (M.S.S.), U19AI109976 (S.C., A.W.G., and M.E.P.), P01AI145815 (S.C., A.W.G., and M.E.P.), and the Frenchman’s Creek Women for Cancer Research (M.E.P.). We thank the The NIH Tetramer Facility which is supported by contract 75N93020D00005. We would like to thank Drs. Mark Sundrud, Kendall Nettles, and John Chang for critically reading the manuscript.

## Author contributions

H.D., M.E.P. and S.C. conceived the study. H.D. and M.E.P. wrote the manuscript with help from A.W.G, S.C., J.J.M., and G.M.. H.D. performed bioinformatics analysis and conducted experiments. R.C. conducted RNAi screen. T.C.V. performed initial analysis of RNAi screen. R.C., D.W., S.T., M.A.F., J.K., P.K., A.J.G, C.T. and J.J.M. performed experiments and analysis. D.M.M. provided insights and reagents related to Chd7.

---

**Figure S1.**
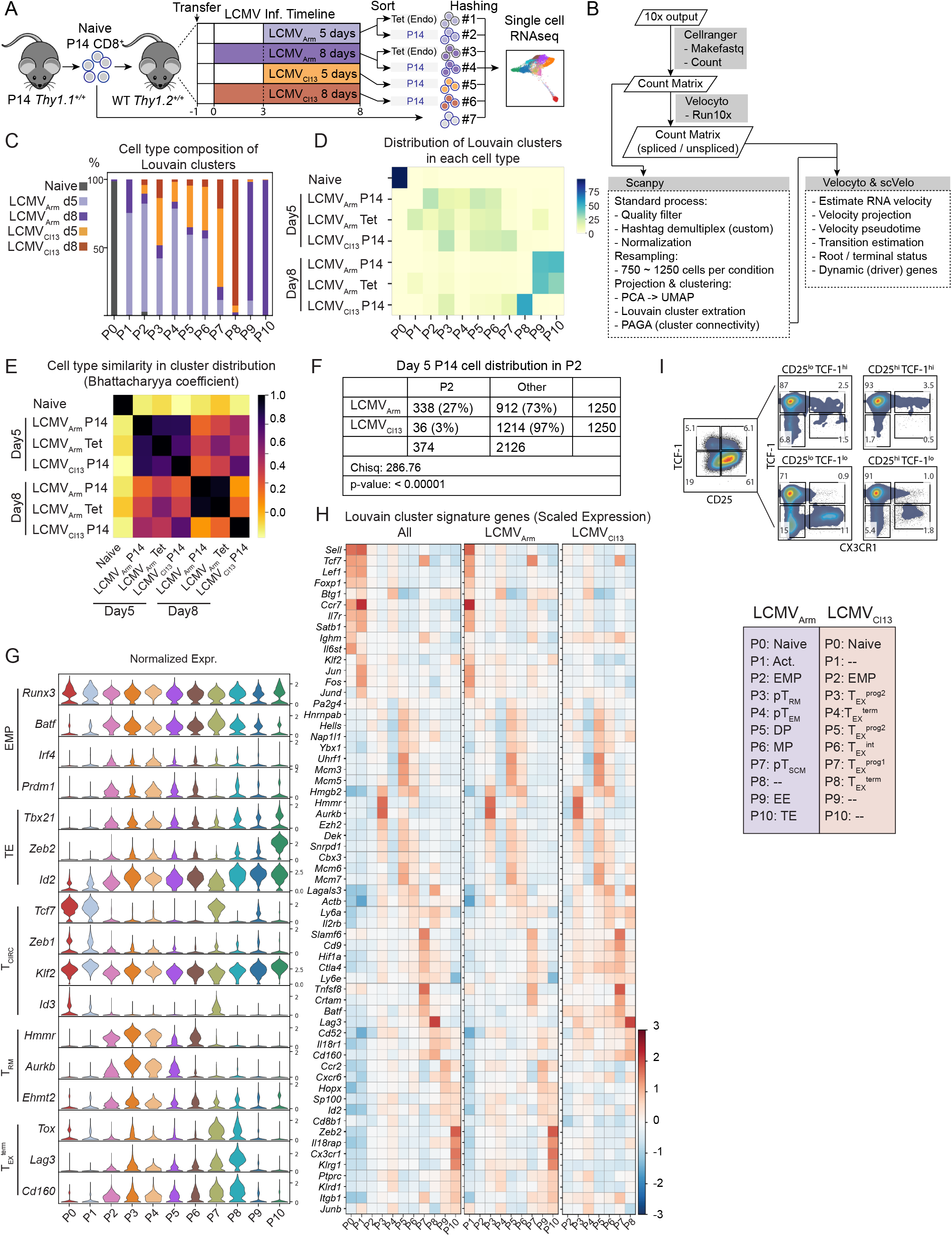
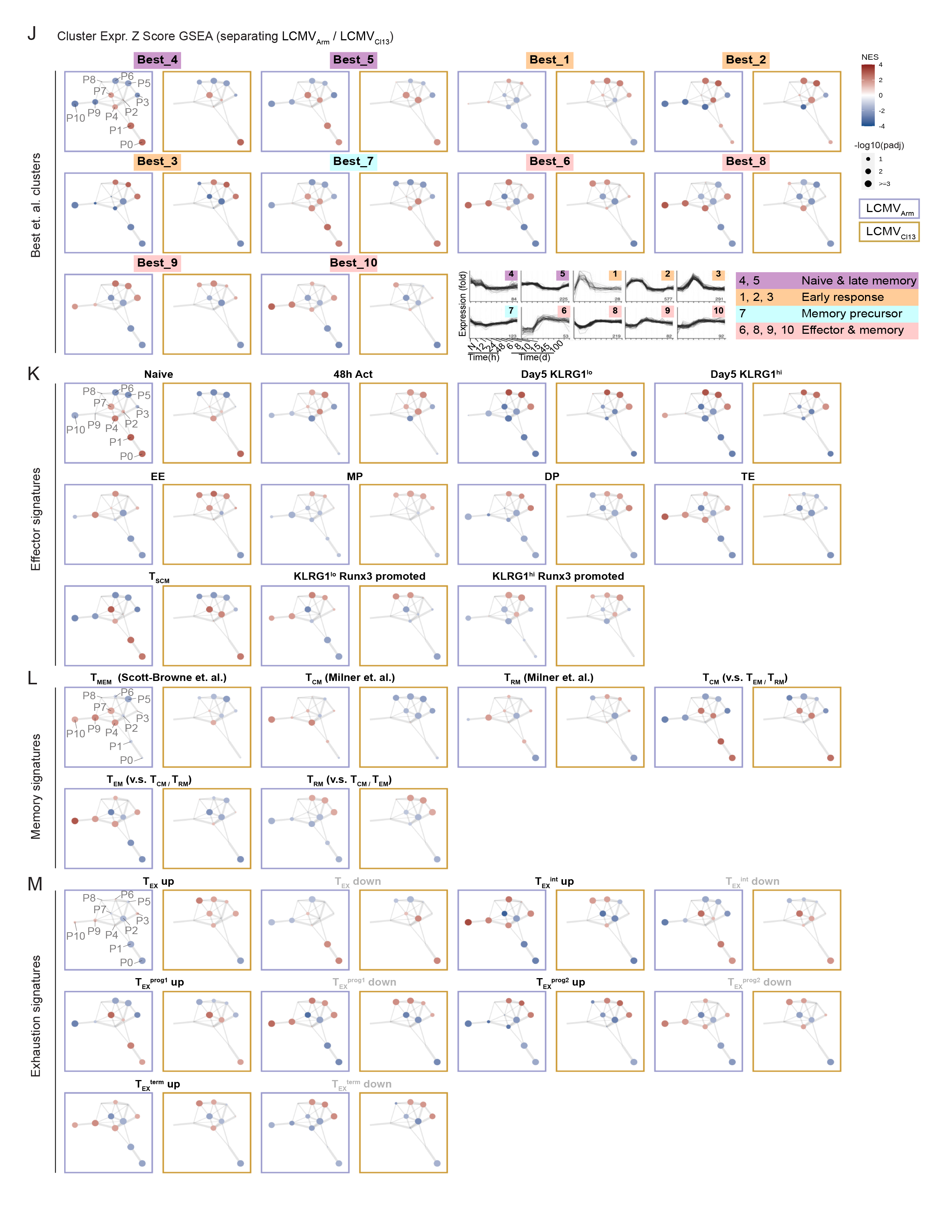
Unsupervised approach to define early developmental clusters of antiviral CD8 T cells during actue and chronic viral infections at single cell level. (A) Schematic of acute v.s. chronic single cell experiment. Donor naive P14 Thy1.1 CD8^+^ T cells from were isolated and transferred into 4 groups of WT recipient mice at day −1. LCMV_Arm_ and LCMV_Cl13_ infections were given at either day 0 or day 3 for each group. P14 Thy1.1 donor CD8^+^ T cells and GP33 Tetramer^+^ endogenous LCMV responding CD8^+^ cells (Tet) were isolated at the same day for each group. Cells of each different origin were hash-tagged and mixed for scRNAseq in the same batch. (B) Schematic of bioinformatic analysis pipeline of acute v.s. chronic single cell experiment. 10x outputs were converted to fastq files and counted for transcript abundance with Cellranger 3.0. Basic quality filter and normalization was performed with Scanpy. Biolegend hashtags were demultiplexed with custom python script. Outliers were detected with DBSCAN (scikit-learn). Cells from each condition were resampled to 750 – 1250 cells condition. Dimentionality reduction (PCA, UMAP, Force Atlas), clustering (Louvain) and cluster connectivity analysis (PAGA) were performed with Scanpy functions. The count matrix was also processed with Velocyto to separate spliced and unspliced transcripts for further velocity associated analysis with scVelo. (C) Stacked bar chart showing composition by different cell type for each Louvain cluster P0-P10. For each cluster, total percentages of all cell types add up to 100%. (D) Heatmap showing percentage distribution in Louvain clusters for each cell type. For each cell type, total percentage in all Louvain clusters add up to 100%. (E) Heatmap represents similarity between cell types based on distribution in Louvain clusters estimated by bhattacharyya coefficient. (F) Chi-square analysis of distribution of day 5 LCMV_Arm_ / LCMV_Cl13_ P14 CD8^+^ cell distribution in / out cluster P2. (G) Violin plots of raw expression (logrithmized and normalized) per cluster for selected phenotype marker genes. (H) See **Fig 1E** description: mean of scaled expression heatmap, top 10 genes ranked by t-test score plotted. Left: all cells; middle and right: separating LCMV_Arm_ / LCMV_Cl13_. (I) Phenotype correlation between CX3CR1 - CD27 and CD25 - TCF-1 of transferred P14 CD8^+^ cell day 4 post LCMV infection (50k cells transferred). (J) (K) (L) (M) GSEA of selected signatures for Louvain clusters, separating LCMV_Arm_ / LCMV_Cl13_. GSEA analysis performed on mean of normalized gene expression per cluster. Colors of dots represent NES, and sizes of dots represent −log10(padj) of signature enrichment. Best et. el. signatures annotated with longitudinal expression dynamics of genes in the clusters and grouped based on expression dynamics (plot from original publication).

**Figure S2.**
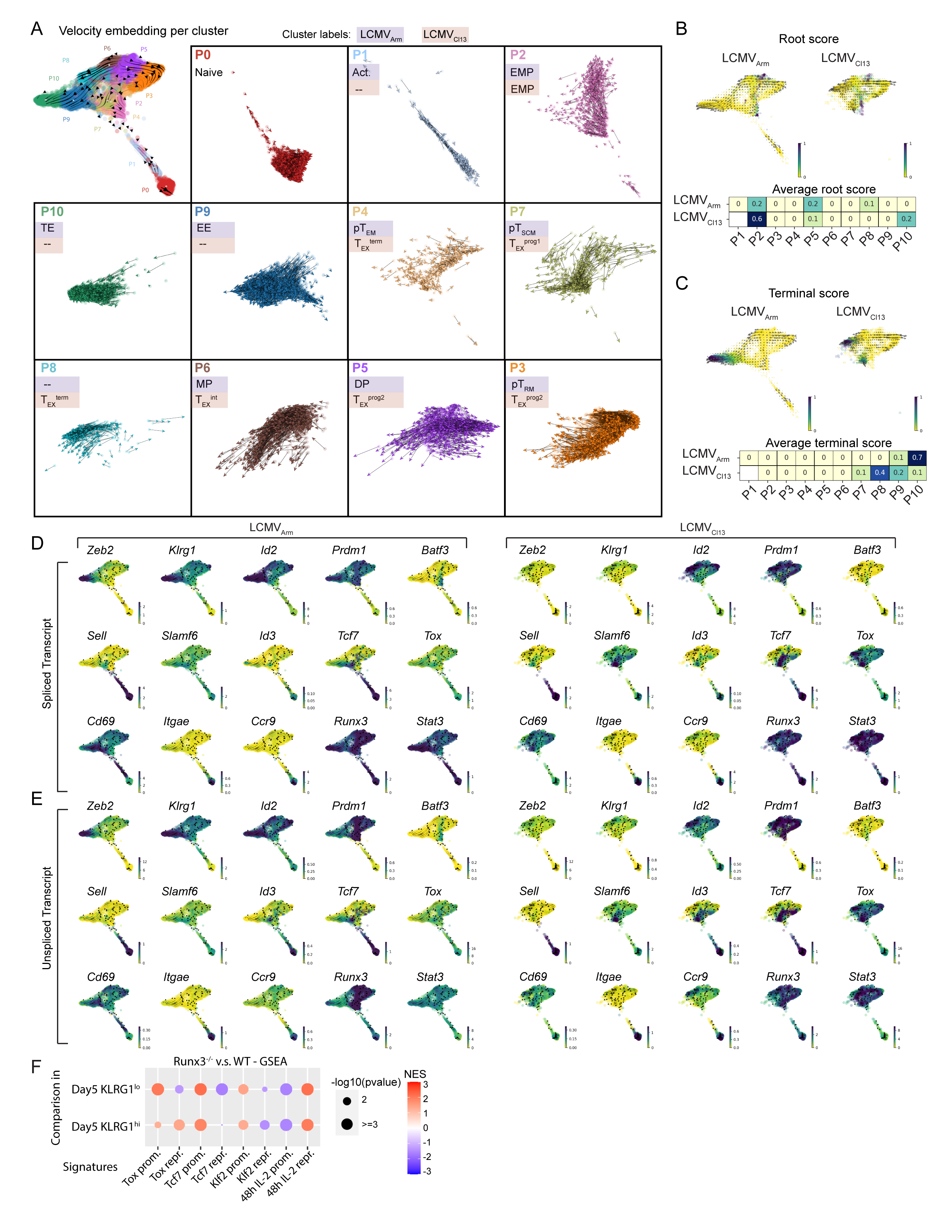
Single cell RNA velocity and root / terminal score indicates P2 & P5 are developmental roots. (A) Single cell velocity vectors for each cluster. (B)(C) Root / terminal score analysis, separating LCMV_Arm_ / LCMV_Cl13_ (not including naive cells). Top: Root / terminal score of single cells, overlayed with grid velocity. Bottom: average root / terminal per cluster. (D)(E) Single cell spliced transcript abundance / unspliced transcript abundance for selected driver genes, Separating LCMV_Arm_ / LCMV_Cl13_. (F) GSEA analysis of P14 Runx3^−/−^ versus WT CD8 T cell gene expression, at day 5 / day 8 p.i. LCMV infection. Positive NES scores indicate signature genes more highly expressed in Runx3^−/−^ comparing with WT; negative NES scores indicate signature genes more highly expressed in WT comparing with Runx3^−/−^.

**Figure S3.**
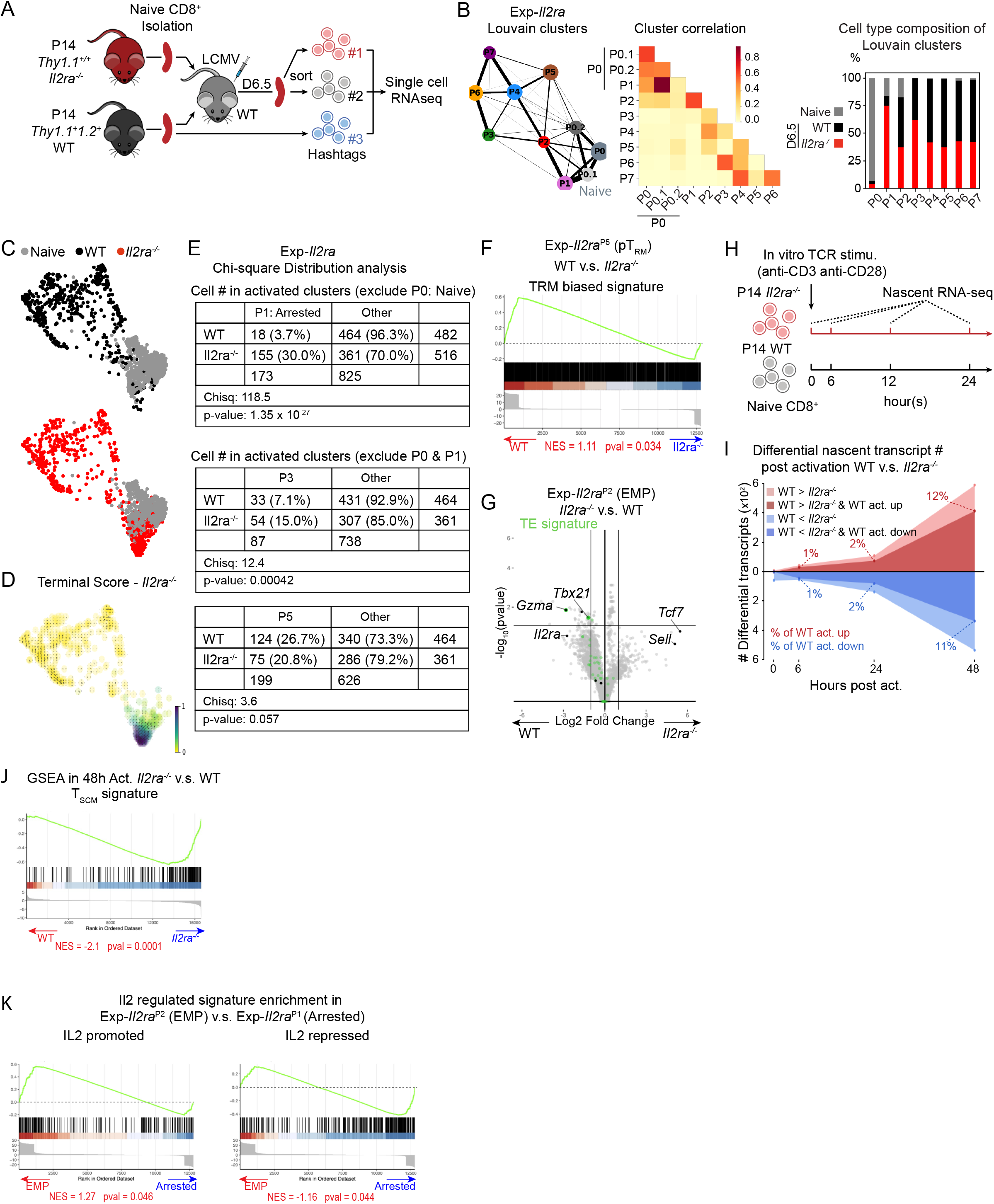
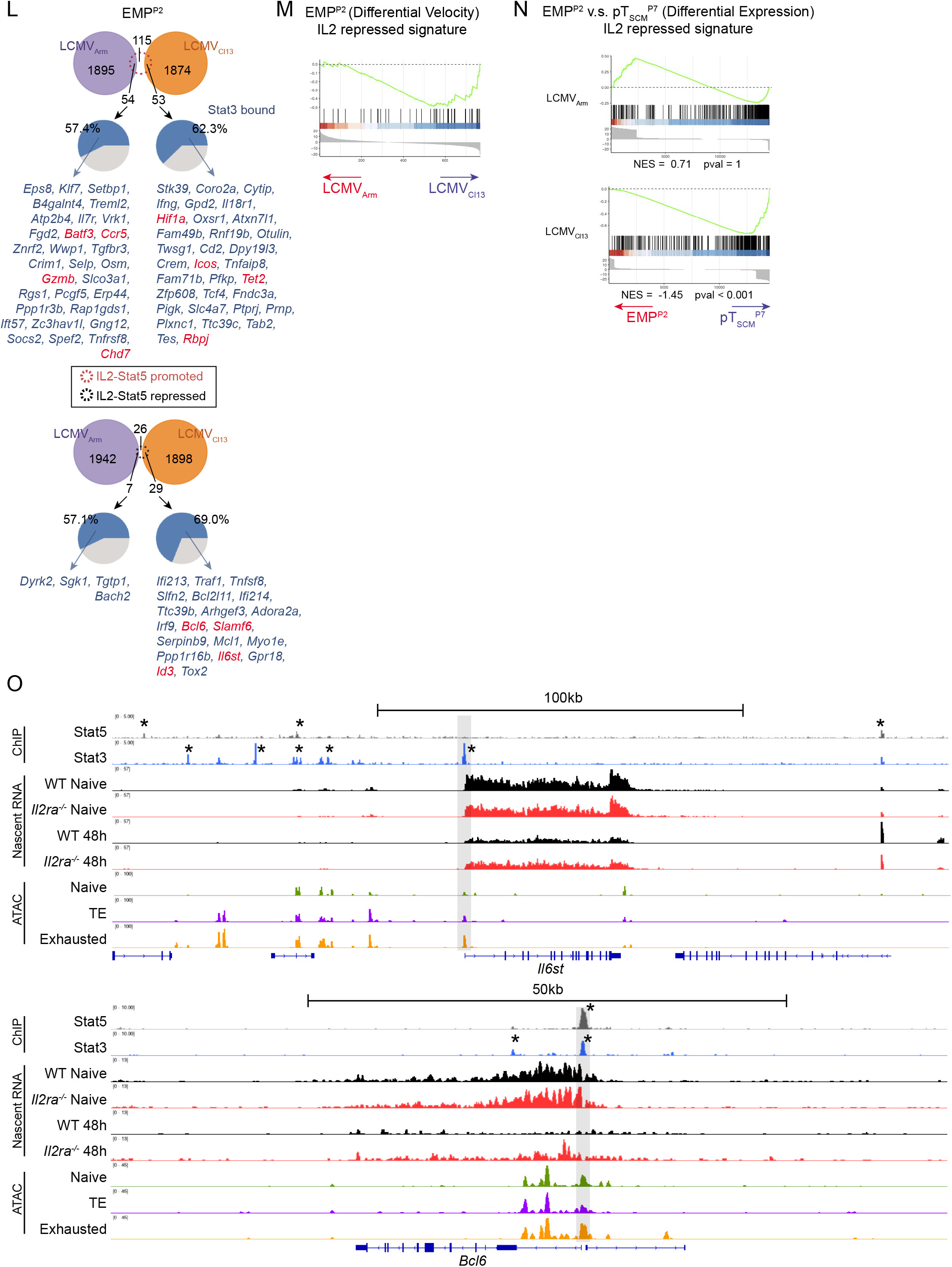
IL-2 signaling contribute to EMP formation via transcription regulation. (A) Schematic of Exp*-Il2ra*: *Il2ra*^-/-^ and WT P14 CD8^+^ T cell cogenetic transfer and single cell experiment. (B) Left: PAGA connectivity graph, each node represent one cluster, node sizes represent relative cell number of cluster, edge widths represent relative PAGA connectivity score. Middle: heatmap of PAGA connectivity score between clusters. Right: Stacked bar chart showing composition by different cell type for each Louvain clusters. For each cluster, total percentages of all cell types add up to 100%. (C) Exp*-Il2ra* PAGA initialized FA embedding, highlighting cells of different origin. (D) Terminal score of *Il2ra^-/-^* single cells in Exp*-Il2ra*. (E) Chi-square analysis of cell type distribution in Exp*-Il2ra*. Top: Cell number distribution in activated clusters in Exp-*Il2ra*^P1^ or outside of Exp-*Il2ra*^P1^ for WT and I *Il2ra^-/-^*. Bottom: Cell number distribution in activated clusters except for P1, comparing WT and *Il2ra*^-/-^. (F) GSEA of T_RM_ gene signature in Exp-*Il2ra*^P5^ WT versus *Il2ra^-/-^*. (G) Differential expression volcano plot of WT and *Il2ra^-/-^* in Exp-*Il2ra*^P2^. The x-axis represents log2 fold change, and y-axis represents −log10(pvalue). The horizontal line indicates pval = 0.05. The vertical lines indicates absolute log2fc = 1. TE signature genes marked in green. (H) Schematic of *Il2ra^-/-^* and WT P14 CD8^+^ T cell *in vitro* activation and nascent RNA-seq experiment. (I) See **Fig 3H**. Dark red and dark blue area represent number of genes genes that are expressed higher in WT than *Il2ra^-/-^* which are upregulated post activation (each time point versus naive), or genes expressed lower in WT than *Il2ra^-/-^* which are downregulated post activation. (J) GSEA of T_SCM_ signature indicates that *Il2ra^-/-^* express T_SCM_ signature genes at higher level than WT at 48h post activation. (K) GSEA enrichment of IL-2 promoted / repressed genes in Exp-*Il2ra*^P3^(EMP) versus Exp-*Il2ra*^P1^ (Arrested) differential gene list (signature genes selected from DEseq2 analysis of 48h post activation, WT versus *Il2ra^-/-^* with cutoffs padj < 0.05, absolute log2 fold change > 1.2). (L) Venn diagrams showing overlap between P2 LCMV_Arm_ and LCMV_Cl13_ differential genes (padj < 0.05) and IL2-Stat5 promoted / repressed genes. Pie charts showing the percentage of Stat3 bound genes in the intersection of LCMV_Arm_ and LCMV_Cl13_ differential genes and IL2-Stat5 promoted / repressed genes. IL2-Stat5 promoted genes: nascent transcript abundance WT > *Il2ra*^-/-^ and WT 48h > WT 6h (DESeq2, pvalue <= 0.05), intersected with genes that are bound by Stat5 ^1^. IL2-Stat5 repressed genes: nascent transcript abundance WT < *Il2ra^-/-^* and WT 48h < WT 6h (DESeq2, pvalue <= 0.05), intersected with genes that are bound by Stat5. (M) GSEA enrichment of IL2 repressed signature between LCMV_Arm_ and LCMV_Cl13_ differential velocity genes in EMP^P2^. (N) GSEA enrichment of 48h IL2 repressed signature (described in fig S3K) in pT_SCM_^P7^ versus EMP^P2^, separating LCMV_Arm_ and LCMV_Cl13_. (O) Visualization of ChIP-seq, ATAC-seq and nascent RNA-seq tracks at *Il6st* and *Bcl6* regions. Asterisks represent significant peaks (MACS2, q-value < 0.01).

**Figure S4.**
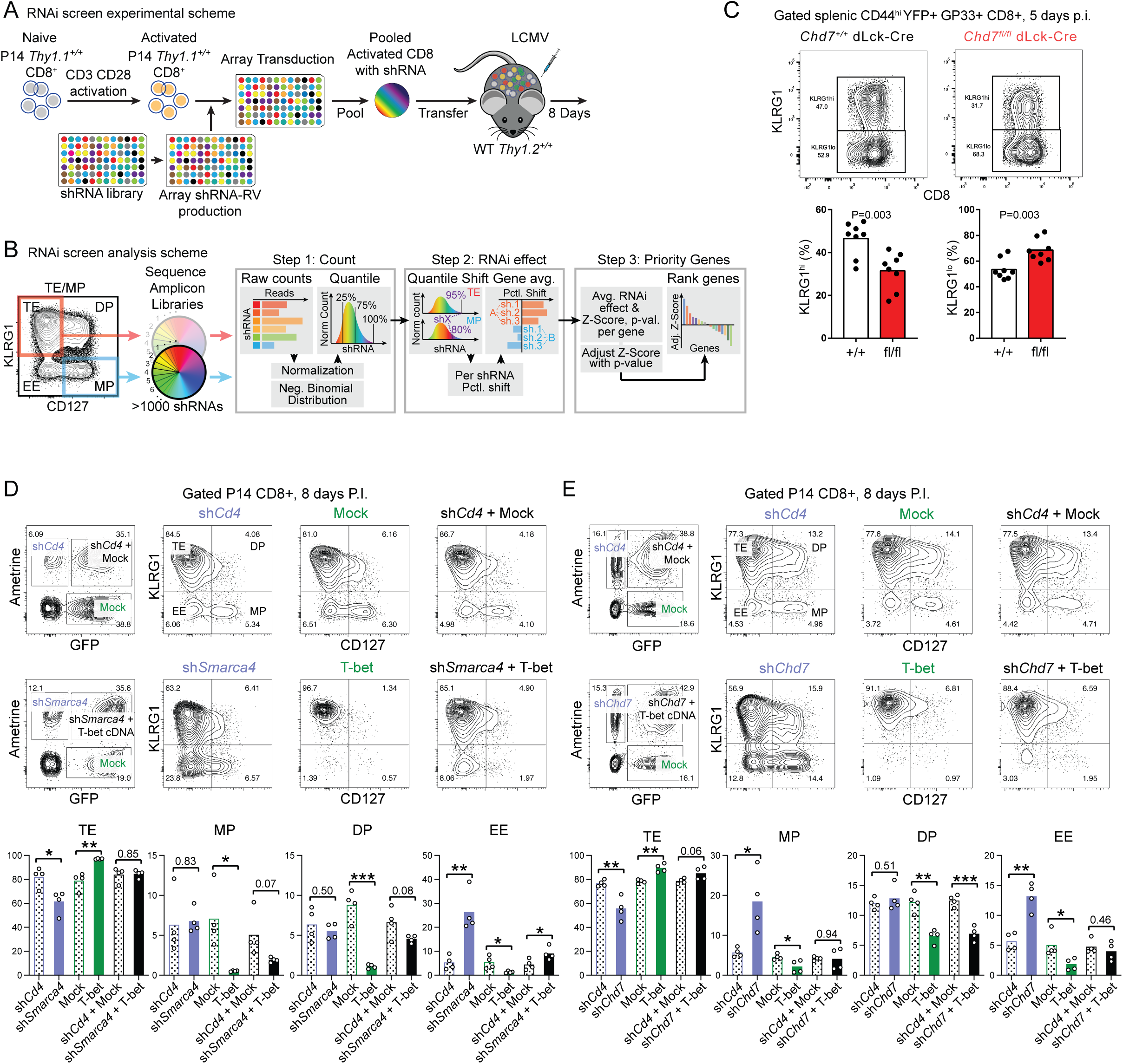
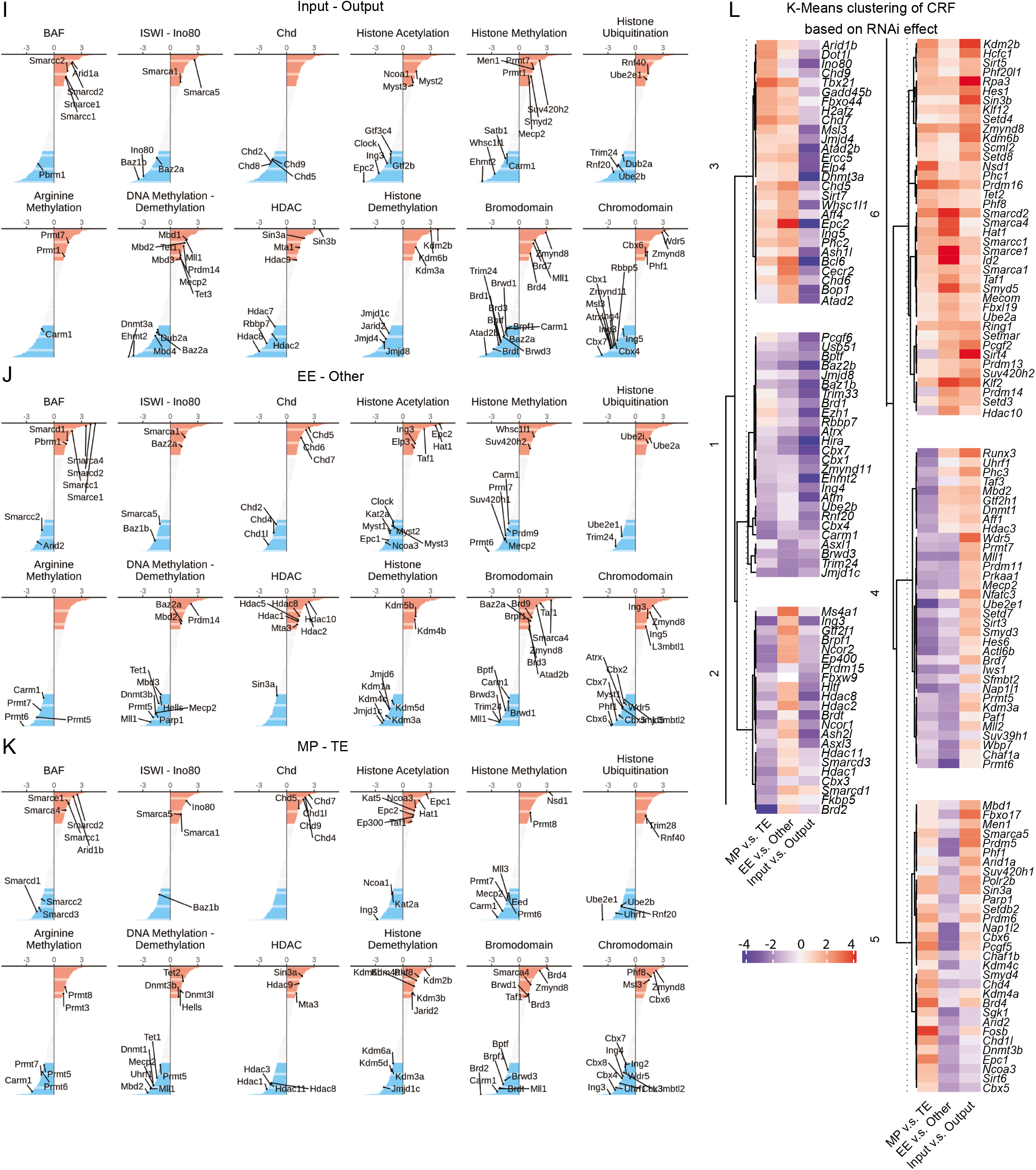
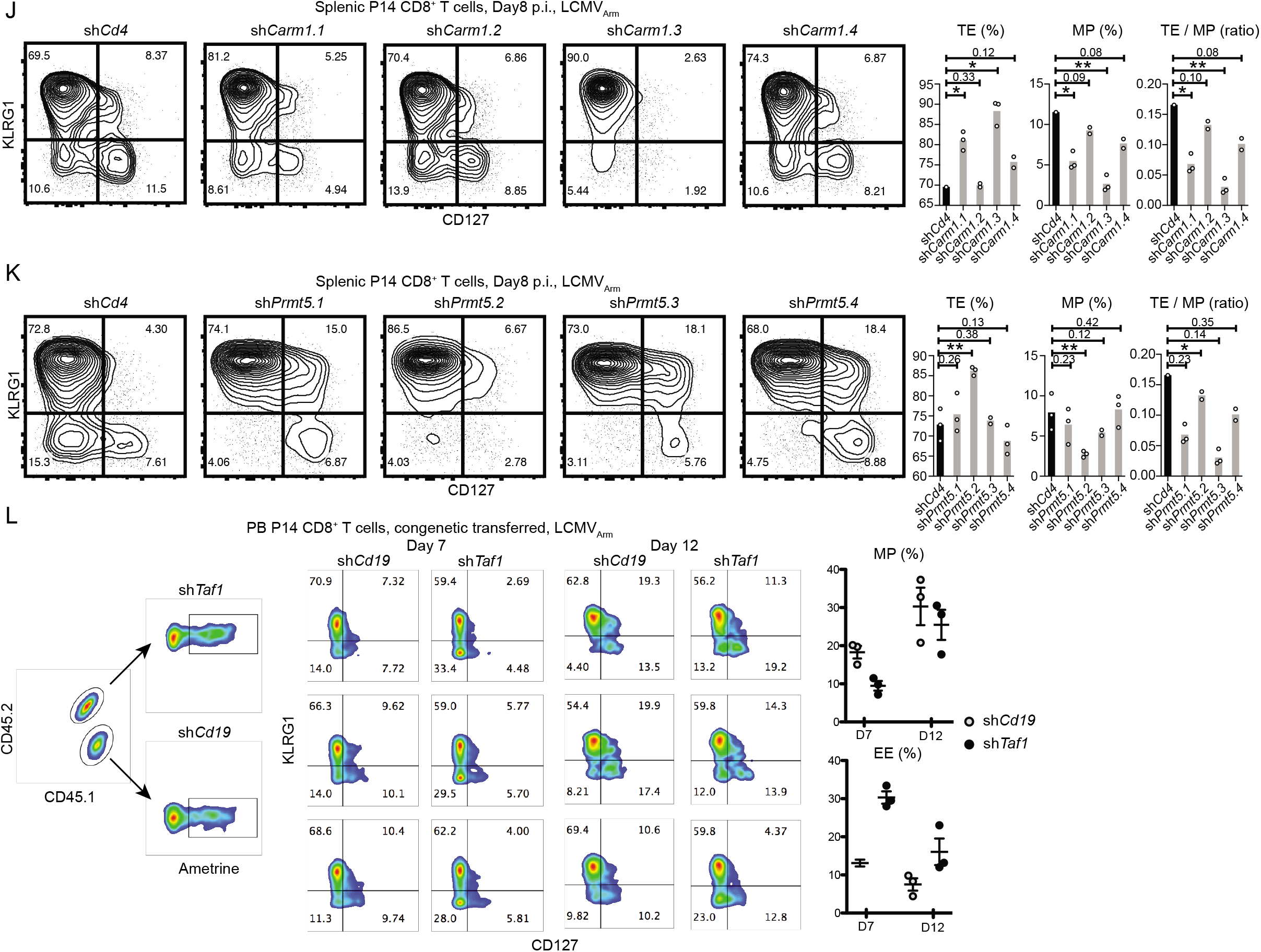
Identification and functional analysis of CRFs that is required for CD8^+^ T cell lineage formation. (A) Experimental setup of CRF RNAi screen (See **Methods**). (B) Bioinformatic analysis pipeline of CRF RNAi screen (See **Methods**). (C) Flow cytometry at day5 pi showing KLRG1^hi^ population in Chd7^fl/fl^ is reduced comparing with Chd7^+/+^. (D)(E) Flow cytometry of transduced and transferred P14 CD8 T cells post LCMV_Arm_ infection. CD8 T cells were simultaneously transduced with a combination of retroviral vectors containing sh*Cd4* (Ametrine, control) and empty vector (GFP, control), or a combination of retroviral vectors containing sh*Smarca4* / sh*Chd7* (Ametrine) and T-bet cDNA (GFP). Phenotypes of RNAi and T-bet overexpression were accessed by flow cytometry at day8 pi. (F)(G)(H) Ranked lists of adjusted RNA - effects for input versus output, MP versus TE, and EE versus others (as described in fig 5B-D), annotating genes from major CRF families. (I) Six major clusters of CRFs extracted from hierarchical clustering of CRFs based on RNAi effect scores of 3 categories (MP v.s. TE, EE v.s. Other, Input v.s. Output) from screen. (J)(K) Flow cytometry of splenic CD8^+^ T cell phenotype day8 pi LCMV_Arm_, comparing transferred P14 CD8 + T cells transduced with sh*Cd4*(control) and multiple shRNAs against Carm1 and Prmt5. (L) Flow cytometry of peripheral blood CD8^+^ T cell phenotype day7 and day12 pi LCMV_Arm_, comparing congenially transferred P14 CD8 + T cells transduced with sh*Cd19*(control) and sh*Taf1*.

**Table S1.** Acute versus Chronic infection (LCMV_Arm_ / LCMV_Cl13_) single cell experiment - clusters and cluster signature genes

**Table S2.** GSEA signatures

**Table S3.** Acute versus Chronic infection (LCMV_Arm_ / LCMV_Cl13_) single cell experiment - differential expression of genes between LCMV_Arm_ and LCMV_Cl13_

**Table S4.** Acute versus Chronic infection (LCMV_Arm_ / LCMV_Cl13_) single cell experiment - scVelo likelihood genes

**Table S5.** Acute versus Chronic infection (LCMV_Arm_ / LCMV_Cl13_) single cell experiment - spliced / unspliced transcript abundance and velocity of likelihood genes

**Table S6.** Exp*-Il2ra* - clusters and cluster signature genes

**Table S7.** RNAi screen of chromatin remodelers: shRNA sequences; scores of each gene in different comparisons

## References

1 Akondy, R. S. et al. Origin and differentiation of human memory CD8 T cells after vaccination. Nature 552, 362–367, doi:10.1038/nature24633 (2017).

2 Jameson, S. C. & Masopust, D. Understanding Subset Diversity in T Cell Memory. Immunity 48, 214–226, doi:10.1016/j.immuni.2018.02.010 (2018).

3 McLane, L. M., Abdel-Hakeem, M. S. & Wherry, E. J. CD8 T Cell Exhaustion During Chronic Viral Infection and Cancer. Annu Rev Immunol 37, 457–495, doi:10.1146/annurev-immunol-041015-055318 (2019).

4 Chung, H. K., McDonald, B. & Kaech, S. M. The architectural design of CD8+ T cell responses in acute and chronic infection: Parallel structures with divergent fates. J Exp Med 218, doi:10.1084/jem.20201730 (2021).

5 Restifo, N. P. & Gattinoni, L. Lineage relationship of effector and memory T cells. Curr Opin Immunol 25, 556–563, doi:10.1016/j.coi.2013.09.003 (2013).

6 Wherry, E. J. et al. Lineage relationship and protective immunity of memory CD8 T cell subsets. Nature immunology 4, 225–234, doi:10.1038/ni889 (2003).

7 Kaech, S. M. & Wherry, E. J. Heterogeneity and cell-fate decisions in effector and memory CD8+ T cell differentiation during viral infection. Immunity 27, 393–405, doi:S1074-7613(07)00410-4 [pii] 10.1016/j.immuni.2007.08.007 (2007).

8 Buchholz, V. R., Schumacher, T. N. & Busch, D. H. T Cell Fate at the Single-Cell Level. Annu Rev Immunol 34, 65–92, doi:10.1146/annurev-immunol-032414-112014 (2016).

9 Reiner, S. L. & Adams, W. C. Lymphocyte fate specification as a deterministic but highly plastic process. Nat Rev Immunol 14, 699–704, doi:10.1038/nri3734 (2014).

10 Rosato, P. C., Wijeyesinghe, S., Stolley, J. M. & Masopust, D. Integrating resident memory into T cell differentiation models. Curr Opin Immunol 63, 35–42, doi:10.1016/j.coi.2020.01.001 (2020).

11 Bergen, V., Lange, M., Peidli, S., Wolf, F. A. & Theis, F. J. Generalizing RNA velocity to transient cell states through dynamical modeling. Nat Biotechnol 38, 1408–1414, doi:10.1038/s41587-020-0591-3 (2020).

12 La Manno, G. et al. RNA velocity of single cells. Nature 560, 494–498, doi:10.1038/s41586-018-0414-6 (2018).

13 Wolf, F. A. et al. PAGA: graph abstraction reconciles clustering with trajectory inference through a topology preserving map of single cells. Genome Biol 20, 59, doi:10.1186/s13059-019-1663-x (2019).

14 Chen, R. et al. *In vivo* RNA interference screens identify regulators of antiviral CD4(+) and CD8(+) T cell differentiation. Immunity 41, 325–338, doi:10.1016/j.immuni.2014.08.002 (2014).

15 Wherry, E. J., Blattman, J. N., Murali-Krishna, K., van der Most, R. & Ahmed, R. Viral persistence alters CD8 T-cell immunodominance and tissue distribution and results in distinct stages of functional impairment. J Virol 77, 4911–4927 (2003).

16 D’Souza, W. N. & Hedrick, S. M. Cutting edge: latecomer CD8 T cells are imprinted with a unique differentiation program. J Immunol 177, 777–781, doi:10.4049/jimmunol.177.2.777 (2006).

17 Blondel, V. D., Guillaume, J. L., Lambiotte, R. & Lefebvre, E. Fast unfolding of communities in large networks. J Stat Mech-Theory E, doi:Artn P10008 10.1088/1742-5468/2008/10/P10008 (2008).

18 Becht, E. et al. Dimensionality reduction for visualizing single-cell data using UMAP. Nat Biotechnol 37, 38–44 (2019).

19 Subramanian, A. et al. Gene set enrichment analysis: a knowledge-based approach for interpreting genome-wide expression profiles. Proc Natl Acad Sci U S A 102, 15545–15550, doi:10.1073/pnas.0506580102 (2005).

20 Yu, G., Wang, L. G., Han, Y. & He, Q. Y. clusterProfiler: an R package for comparing biological themes among gene clusters. OMICS 16, 284–287, doi:10.1089/omi.2011.0118 (2012).

21 Best, J. A. et al. Transcriptional insights into the CD8(+) T cell response to infection and memory T cell formation. Nature immunology 14, 404–412, doi:10.1038/ni.2536 (2013).

22 Stauber, D. J., Debler, E. W., Horton, P. A., Smith, K. A. & Wilson, I. A. Crystal structure of the IL-2 signaling complex: paradigm for a heterotrimeric cytokine receptor. Proc Natl Acad Sci U S A 103, 2788–2793, doi:10.1073/pnas.0511161103 (2006).

23 Wang, X., Rickert, M. & Garcia, K. C. Structure of the quaternary complex of interleukin-2 with its alpha, beta, and gammac receptors. Science 310, 1159–1163, doi:10.1126/science.1117893 (2005).

24 Diao, H. & Pipkin, M. Stability and flexibility in chromatin structure and transcription underlies memory CD8 T-cell differentiation. F1000Res 8, doi:10.12688/f1000research.18211.1 (2019).

25 Wang, D. et al. The Transcription Factor Runx3 Establishes Chromatin Accessibility of cis-Regulatory Landscapes that Drive Memory Cytotoxic T Lymphocyte Formation. Immunity 48, 659–674 e656, doi:10.1016/j.immuni.2018.03.028 (2018).

26 Kaech, S. M. & Cui, W. Transcriptional control of effector and memory CD8+ T cell differentiation. Nature reviews. Immunology 12, 749–761, doi:10.1038/nri3307 (2012).

27 Milner, J. J. & Goldrath, A. W. Transcriptional programming of tissue-resident memory CD8(+) T cells. Curr Opin Immunol 51, 162–169, doi:10.1016/j.coi.2018.03.017 (2018).

28 Shu, J. et al. Induction of Pluripotency in Mouse Somatic Cells with Lineage Specifiers. Cell 153, 963–975, doi:10.1016/j.cell.2013.05.001 (2013).

29 Laslo, P. et al. Multilineage transcriptional priming and determination of alternate hematopoietic cell fates. Cell 126, 755–766, doi:10.1016/j.cell.2006.06.052 (2006).

30 Joshi, N. S. et al. Inflammation directs memory precursor and short-lived effector CD8(+) T cell fates via the graded expression of T-bet transcription factor. Immunity 27, 281–295, doi:10.1016/j.immuni.2007.07.010 (2007).

31 Dominguez, C. X. et al. The transcription factors ZEB2 and T-bet cooperate to program cytotoxic T cell terminal differentiation in response to LCMV viral infection. J Exp Med 212, 2041–2056, doi:10.1084/jem.20150186 (2015).

32 Omilusik, K. D. et al. Transcriptional repressor ZEB2 promotes terminal differentiation of CD8+ effector and memory T cell populations during infection. J Exp Med 212, 2027–2039, doi:10.1084/jem.20150194 (2015).

33 Beltra, J. C. et al. Developmental Relationships of Four Exhausted CD8(+) T Cell Subsets Reveals Underlying Transcriptional and Epigenetic Landscape Control Mechanisms. Immunity 52, 825–841 e828, doi:10.1016/j.immuni.2020.04.014 (2020).

34 Milner, J. J. et al. Runx3 programs CD8(+) T cell residency in non-lymphoid tissues and tumours. Nature 552, 253–257, doi:10.1038/nature24993 (2017).

35 Alfei, F. et al. TOX reinforces the phenotype and longevity of exhausted T cells in chronic viral infection. Nature 571, 265–269, doi:10.1038/s41586-019-1326-9 (2019).

36 Khan, O. et al. TOX transcriptionally and epigenetically programs CD8(+) T cell exhaustion. Nature 571, 211–218, doi:10.1038/s41586-019-1325-x (2019).

37 Yao, C. et al. Single-cell RNA-seq reveals TOX as a key regulator of CD8(+) T cell persistence in chronic infection. Nat Immunol 20, 890–901, doi:10.1038/s41590-019-0403-4 (2019).

38 Pipkin, M. E. Runx proteins and transcriptional mechanisms that govern memory CD8 T cell development. Immunol Rev 300, 100–124, doi:10.1111/imr.12954 (2021).

39 Wu, J. I., Lessard, J. & Crabtree, G. R. Understanding the words of chromatin regulation. Cell 136, 200–206, doi:10.1016/j.cell.2009.01.009 (2009).

40 Bajpai, R. et al. CHD7 cooperates with PBAF to control multipotent neural crest formation. Nature 463, 958–962, doi:10.1038/nature08733 (2010).

41 Gennery, A. R. et al. Mutations in CHD7 in patients with CHARGE syndrome cause T-B + natural killer cell + severe combined immune deficiency and may cause Omenn-like syndrome. Clin Exp Immunol 153, 75–80, doi:10.1111/j.1365-2249.2008.03681.x (2008).

42 Hurd, E. A., Poucher, H. K., Cheng, K., Raphael, Y. & Martin, D. M. The ATP-dependent chromatin remodeling enzyme CHD7 regulates pro-neural gene expression and neurogenesis in the inner ear. Development 137, 3139–3150, doi:10.1242/dev.047894 (2010).

43 Writzl, K., Cale, C. M., Pierce, C. M., Wilson, L. C. & Hennekam, R. C. Immunological abnormalities in CHARGE syndrome. Eur J Med Genet 50, 338–345, doi:10.1016/j.ejmg.2007.05.002 (2007).

44 Chi, T. H. et al. Reciprocal regulation of CD4/CD8 expression by SWI/SNF-like BAF complexes. Nature 418, 195–199, doi:10.1038/nature00876 (2002).

45 Mackay, L. K. et al. Hobit and Blimp1 instruct a universal transcriptional program of tissue residency in lymphocytes. Science 352, 459–463, doi:10.1126/science.aad2035 (2016).

46 Xin, A. et al. A molecular threshold for effector CD8(+) T cell differentiation controlled by transcription factors Blimp-1 and T-bet. Nat Immunol 17, 422–432, doi:10.1038/ni.3410 (2016).

47 Wu, T. et al. The TCF1-Bcl6 axis counteracts type I interferon to repress exhaustion and maintain T cell stemness. Science immunology 1 (2016).

48 Keppler, S. J., Rosenits, K., Koegl, T., Vucikuja, S. & Aichele, P. Signal 3 cytokines as modulators of primary immune responses during infections: the interplay of type I IFN and IL-12 in CD8 T cell responses. PLoS One 7, e40865, doi:10.1371/journal.pone.0040865 (2012).

49 Kurd, N. S. et al. Early precursors and molecular determinants of tissue-resident memory CD8(+) T lymphocytes revealed by single-cell RNA sequencing. Sci Immunol 5, doi:10.1126/sciimmunol.aaz6894 (2020).

50 Milner, J. J. et al. Heterogenous Populations of Tissue-Resident CD8(+) T Cells Are Generated in Response to Infection and Malignancy. Immunity 52, 808–824 e807, doi:10.1016/j.immuni.2020.04.007 (2020).

51 Fonseca, R. et al. Developmental plasticity allows outside-in immune responses by resident memory T cells. Nat Immunol 21, 412–421, doi:10.1038/s41590-020-0607-7 (2020).

52 Youngblood, B. et al. Effector CD8 T cells dedifferentiate into long-lived memory cells. Nature 552, 404–409, doi:10.1038/nature25144 (2017).

53 Pauken, K. E. et al. Epigenetic stability of exhausted T cells limits durability of reinvigoration by PD-1 blockade. Science 354, 1160–1165, doi:10.1126/science.aaf2807 (2016).

54 Sen, D. R. et al. The epigenetic landscape of T cell exhaustion. Science 354, 1165–1169, doi:10.1126/science.aae0491 (2016).

55 Li, P. et al. STAT5-mediated chromatin interactions in superenhancers activate IL-2 highly inducible genes: Functional dissection of the Il2ra gene locus. Proceedings of the National Academy of Sciences 114, 12111–12119 (2017).

56 Rodda, L. B. et al. Single-Cell RNA Sequencing of Lymph Node Stromal Cells Reveals Niche-Associated Heterogeneity. Immunity 48, 1014–+, doi:10.1016/j.immuni.2018.04.006 (2018).

57 Langmead, B. & Salzberg, S. L. Fast gapped-read alignment with Bowtie 2. Nat Methods 9, 357–359 (2012).

58 Li, H. et al. The sequence alignment/map format and SAMtools. Bioinformatics 25, 2078–2079 (2009).

59 Liao, Y., Smyth, G. K. & Shi, W. The Subread aligner: fast, accurate and scalable read mapping by seed- and-vote. Nucleic acids research 41, e108–e108 (2013).

60 Love, M. I., Huber, W. & Anders, S. Moderated estimation of fold change and dispersion for RNA-seq data with DESeq2. Genome Biol 15, 550, doi:10.1186/s13059-014-0550-8 (2014).

61 Wolf, F. A., Angerer, P. & Theis, F. J. SCANPY: large-scale single-cell gene expression data analysis. Genome Biol 19, 15, doi:10.1186/s13059-017-1382-0 (2018).

62 Korsunsky, I. et al. Fast, sensitive and accurate integration of single-cell data with Harmony. Nat Methods 16, 1289–1296 (2019).

63 Amemiya, H. M., Kundaje, A. & Boyle, A. P. The ENCODE blacklist: identification of problematic regions of the genome. Scientific reports 9, 1–5 (2019).

64 Quinlan, A. R. & Hall, I. M. BEDTools: a flexible suite of utilities for comparing genomic features. Bioinformatics 26, 841–842 (2010).

65 Zhang, Y. et al. Model-based analysis of ChIP-Seq (MACS). Genome Biol 9, R137, doi:10.1186/gb-2008-9-9-r137 (2008).

66 Yu, G., Wang, L. G. & He, Q. Y. ChIPseeker: an R/Bioconductor package for ChIP peak annotation, comparison and visualization. Bioinformatics 31, 2382–2383, doi:10.1093/bioinformatics/btv145 (2015).

